# Transcriptomic landscape and potential therapeutic targets for human testicular aging revealed by single-cell RNA sequencing

**DOI:** 10.1101/2022.12.11.519976

**Authors:** Kai Xia, Siyuan He, Peng Luo, Lin Dong, Feng Gao, Xuren Chen, Yunlin Ye, Yong Gao, Yuanchen Ma, Yadong Zhang, Qiyun Yang, Dayu Han, Xin Feng, Zi Wan, Hongcai Cai, Qiong Ke, Tao Wang, Weiqiang Li, Xiang’an Tu, Xiangzhou Sun, Chunhua Deng, Andy Peng Xiang

## Abstract

**Background:** Testicular aging is known to cause male age-related fertility decline and hypogonadism, but the underlying molecular mechanisms remain unclear.

**Methods:** We survey the single-cell transcriptomic landscape of testes from young and old men and examine age-related changes in germline and somatic niche cells.

**Results:** In-depth evaluation of the gene expression dynamics of germline cells reveals that disturbance of base-excision repair pathway is a major feature of aging spermatogonial stem cells (SSCs), suggesting that defective DNA repair of SSCs may serve as a potential driver for increased de novo germline mutations with age. Further analysis of aging-associated transcriptional changes shows that stress-related changes and apoptotic signaling pathway accumulate in aged somatic cells. We identify age-related impairment of redox homeostasis in aged Leydig cells and find that pharmacological treatment with antioxidants alleviate this cellular dysfunction of Leydig cells and promote testosterone production. Lastly, our results reveal that decreased pleiotrophin (PTN) signaling is a contributing factor for testicular aging.

**Conclusions:** These findings provide a comprehensive understanding of the cell-type-specific mechanisms underlying human testicular aging at a single-cell resolution, and suggest potential therapeutic targets that may be leveraged to address age-related male fertility decline and hypogonadism.

**Funding:** This work was supported by the National Key Research and Development Program of China (2018YFA0107200, 2018YFA0801404), the National Natural Science Foundation of China (32130046, 82171564, 82101669, 81871110, 81971759), the Key Research and Development Program of Guangdong Province (2019B020234001), the Natural Science Foundation of Guangdong Province, China (2022A1515010371), the Major Project of Medical Science and Technology Development Research Center of National Health Planning Commission, China (HDSL202001000), the Open Project of NHC Key Laboratory of Male Reproduction and Genetics (Family Planning Research Institute of Guangdong Province) (KF202001), the Guangdong Province Regional Joint Fund-Youth Fund Project (2021A1515110921), the China Postdoctoral Science Foundation (2021M703736).

## Introduction

The testis is a critical male reproductive organ that serves as the source of sperm and a major supplier of the sex hormone, testosterone(Makela, Koskenniemi, Virtanen, & Toppari, 2019). Thus, the testis is indispensable for both male fertility maintenance and endocrine homeostasis. However, testicular function declines gradually as men age. Previous studies have shown that aging negatively affects sperm parameters, sperm DNA integrity, genomic DNA mutations, chromosomal structures, and epigenetic factors(S. L. Johnson, Dunleavy, Gemmell, & Nakagawa, 2015; Matzkin, Calandra, Rossi, Bartke, & Frungieri, 2021). Despite this, it is becoming more common for men in developed countries to first become a parent at an older age, raising concerns about declines in fertilizing capacity and the consequences for the offspring’s health(Laurentino et al., 2020). Aging also impairs testosterone production and causes male hypogonadism, which is characterized by low libido, erectile dysfunction, infertility, obesity, muscle weakness, osteoporosis, depressed mood, impaired cognition, and other symptoms(Kaufman, Lapauw, Mahmoud, T’Sjoen, & Huhtaniemi, 2019; Mularoni et al., 2020). Testicular aging therefore affects not only men’s reproductive functions, but also their overall health status and quality of life(Matzkin et al., 2021). Consequently, it is important to elucidate the mechanisms underlying testicular aging and identify interventions that might slow or postpone this process.

The testis is a complex structure consisting of numerous heterogeneous cell types, including germline cells at different developmental stages and several somatic cell types(Guo et al., 2018). The propagation of the male germline depends on a specialized cell population called spermatogonial stem cells (SSCs)(Sharma, Wistuba, Pock, Schlatt, & Neuhaus, 2019). SSCs have unlimited self-renewal capacity and, when induced to differentiate, give rise to a series of germ cell stages that ultimately generate sperm in a highly orchestrated manner within the seminiferous tubules(Sharma et al., 2019). The testis niche plays an important role in regulating the survival and differentiation of the male germline(Oatley & Brinster, 2012). In the adult testis, somatic niche cells, including Sertoli cells (SCs)(S. R. Chen & Liu, 2015), Leydig cells (LCs)(Oatley, Oatley, Avarbock, Tobias, & Brinster, 2009), peritubular myoid cells (PTMs)(L. Y. Chen, Brown, Willis, & Eddy, 2014), etc., provide physical and hormonal support for successful spermatogenesis from SSCs. In particular, LCs are responsible for the biosynthesis of testosterone, which acts on target cells in the testes and elsewhere to promote spermatogenesis and male-associated characteristics(Zirkin & Papadopoulos, 2018). Previous studies have shown that aging testes undergo profound morphological alterations of germ cells and somatic cells, leading to reduced functionality(Jiang et al., 2014; Santiago, Silva, Alves, Oliveira, & Fardilha, 2019). However, the cellular and molecular alterations that underlie these changes have not been systematically explored and remain largely unknown.

For a heterogeneous organ such as the testis, it is difficult to accurately reveal cell-type-specific changes in gene expression using conventional bulk RNA-sequencing (RNA-seq) approaches. With advances in the single-cell RNA sequencing (scRNA-seq) technique, it is now possible to analyze alterations of gene transcription within highly heterogeneous tissues at the single-cell level(Di Persio et al., 2021; Guo et al., 2018). Recently, several scRNA-seq studies have begun to lift the veil on the full compendium of gene expression phenotypes and changes in spermatogenic and somatic cells, demonstrating the power of scRNA-seq profiling as a means to study human testis at the single-cell resolution(Guo et al., 2018; Guo et al., 2020; Mahyari et al., 2021). These datasets have revealed the previously obscured molecular heterogeneity among and between varied testicular cell types and are reinvigorating the investigation of testicular biology and pathology(Alfano et al., 2021; Di Persio et al., 2021; Mahyari et al., 2021; Matzkin et al., 2021; Nie et al., 2022; L. Zhao et al., 2020). However, the effect of aging on various testicular cell types in humans has not been analyzed in depth and the critical molecular drivers underlying testicular cell functional decline during aging remain unclear.

In this study, we used human samples to comprehensively survey the single-cell transcriptomic landscape of human testicular aging. We used scRNA-seq to identify gene expression signatures for five germline cell types and six somatic cell types. We examined the transcriptional changes within each major testicular cell type during aging, and report a set of molecular mechanisms underlying human testis aging. Our analysis of aging-associated gene expression changes revealed that DNA repair is disturbed in aged SSCs, and this represents an essential factor underlying the age-related increase in de novo germline mutations. Analysis of human LCs uncovered aging-associated upregulation of oxidative stress response genes, and further experiments revealed that antioxidant treatment alleviated the cellular dysfunction of LCs and promoted testosterone production. An analysis of cell interactions revealed that the pleiotrophin (PTN) signaling pathway is markedly interrupted in aging testis. These data may be used at both bench and bedside in future efforts to address age-related male fertility decline and hypogonadism.

## Results

### Construction of a single-cell transcriptomic atlas of the human testis

To investigate the cellular and molecular alterations of testicular cells during aging, we obtained human testis biopsies from three young individuals (24, 28, and 31 years old) and three aged individuals (61, 70, and 87 years old) (Figure S1A). Histologically, the area occupied by seminiferous tubules was significantly decreased in aged men, compared with young men (Figure 1A). In addition, age-related thickening of boundary tissue was typically found in old men (Figure 1B). These data are in agreement with previous observations(L. Johnson, Abdo, Petty, & Neaves, 1988; Mularoni et al., 2020) and confirmed that this cohort of tissue donors could be used to examine age-related changes in testicular function.

**Figure 1.**
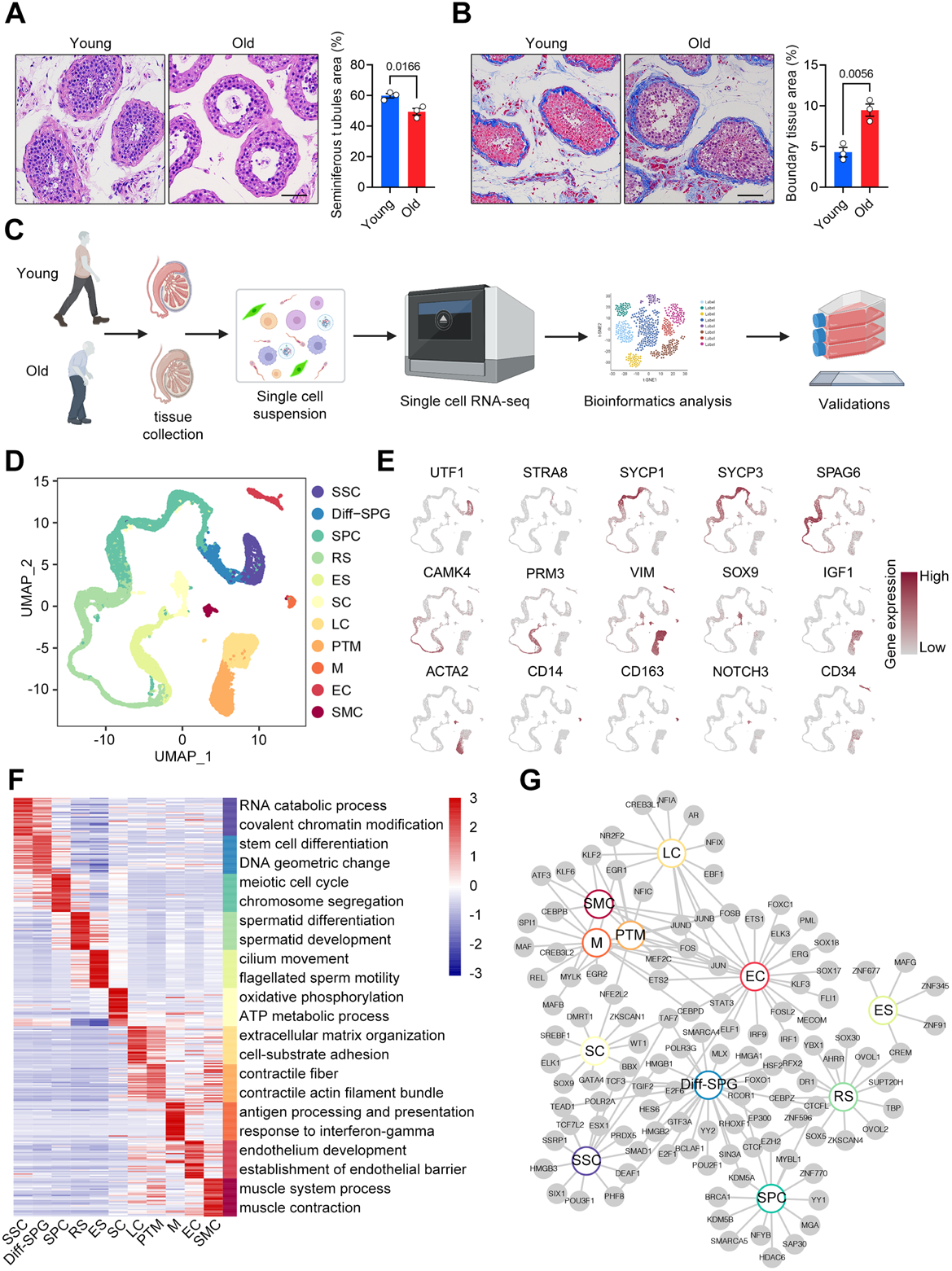
Construction of a Single-Cell Transcriptomic Atlas of Human Testis. (A) Left: representative light micrographs of young and old human testis sections stained with H&E. Right: quantification of the average area of seminiferous tubules. Scale bar, 75 μm. Young, n=3; Old, n=3. Data are expressed as mean ± sem. Significance was determined by Student’s t-test. (B) Left: representative light micrographs of Masson’s trichrome staining of young and old human testis. Right: quantification of the average area of boundary tissue. Scale bar, 75 μm. Young, n=3; Old, n=3. Data are expressed as mean ± sem. Significance was determined by Student’s t-test. (C) Flow chart of scRNA-seq and bioinformatics analysis of the human testis. (D) UMAP plot showing distribution of 11 different cell types in the human testis, with annotation. (E) UMAP plot showing the expression profiles of the indicated cell-type-specific marker genes for the assessed cell types in the human testis. The color key, ranging from gray to brown, indicates low to high gene expression levels, respectively. (F) Heatmap showing the expression profiles of the top 30 cell-type-specific marker genes of different cell types, with their enriched functional annotations on the right. The value for each gene represents scaled data. (G) Network plot showing transcriptional regulators of cell-type-specific marker genes (*p*-value < 0.05, |logFC| > 0.4, min.pct = 0.5) for different cell types in the human testis.

To resolve cell-type-specific alterations in gene expression during testicular aging at the single-cell level, we isolated single cells from these tissues and performed scRNA-seq using the 10x Genomics platform (Figure 1C). We removed cells likely to be of low quality due to debris and doublets, and applied other constraints. Data from 11,444 and 11,076 cells passed quality control and were included in the subsequent analysis for young and old samples, respectively (Figure S1A, S1B). We visualized global human testicular cell populations using uniform manifold approximation and projection (UMAP) and used 27 published marker genes(Alfano et al., 2021; Guo et al., 2018; Guo et al., 2021) to identify the following 11 cell types based on the expression of specific marker genes: SSCs (UTF1+, ID4+), differentiating spermatogonia (Diff-SPGs; STRA8+, DMRT1+), spermatocytes (SPCs; SYCP1+, SYCP3+, and DAZL+), round spermatids (RSs; SPAG6+, CAMK4+), elongating spermatids (ESs; PRM3+, HOOK1+, and TNP1+), SCs (SOX9+, AMH+, and WT1+), LCs (IGF1+, CFD+, and FUM+), PTMs (ACTA2+, MYH11+, NOTCH3-), macrophages (Ms; CD14+, CD163+, and C1QA+), endothelial cells (ECs; CD34+, NOSTRIN+), and smooth muscle cells (SMCs; ACTA2+, MYH11+, NOTCH3+) (Figure 1D,E; Figure S1C-E; Supplementary file 1). Analysis of the top 30 marker genes revealed that each cell type had unique transcriptional features and enriched pathways relevant to their distinct biological functions (Figure 1F; Supplementary file 1). For example, the Gene Ontology (GO) terms “meiosis” and “myofiber” were enriched for SPCs and PTMs, respectively (Figure 1F).

We further constructed a regulatory network of transcription factors (TFs) that defined core TFs unique to each cell type, and hub TFs shared by at least two cell types (Figure 1G; Supplementary file 1). Within the cell type-specific TF networks, the prominent genes included TCF7L2 for SSCs, SOX9 for SCs, and CREB3L1 for LCs. TCF3 was identified in the networks of both SSCs and Diff-SPGs, suggesting that TCF3 plays essential roles in spermatogonia maintenance and specification. FOS was identified in LCs, PTMs, Ms, ECs, and SMCs, indicating that it functions as a broad-acting transcriptional regulator (Figure 1G). This network analysis provided a depiction of the unique and coordinated transcriptional regulatory processes involved in establishing human testicular cell identities.

We next sought to explore cell type-specific gene expression alterations associated with aging. We found similar transcriptional signatures of marker genes for each cell type between young and old humans (Figure S2A), demonstrating that aging had minimal effect on cell identity. In addition to these classic markers, we identified a set of novel markers of testicular cells: SIX1 and LIN7B for SSCs, BEND for SPCs, ABHD4 and TCEAL5 for SCs, KRT17 for PTMs, ADAP2 for Ms, PCAT19 and LCN6 for ECs, and RRAD and RERGL for SMCs (Figure S2B).

Our analyses indicated that the developed atlas could be used to comprehensively delineate the cellular and molecular alterations induced by aging in the human testis.

### Aging-related cellular alterations along the trajectories of spermatogenesis

To identify germline changes during aging, we performed a focused analysis of germ cell clusters. We first investigated the proportions of germ cells at each stage. We identified the following five broad developmental stages of germ cells based on their marker genes: SSCs (UTF1+, ID4+), Diff-SPGs (STRA8+, DMRT1+), SPCs (SYCP1+, SYCP3+, and DAZL+), RSs (SPAG6+, CAMK4+), and ESs (PRM3+, HOOK1+, and TNP1+) (Figures 2A, 2B). We further inferred the trajectories of spermatogenesis by performing an orthogonal pseudotime analysis using the Monocle package(Qiu et al., 2017). This analysis revealed that complete spermatogenesis was present in all samples (Figure 2C, 2D). We then parsed out the germ cells by developmental stage to examine their relative compositions at different ages. Interestingly, these percentages were not significantly different from SSCs to ESs. In contrast, the proportions of germ cells at RS stage and beyond tended to decline with age (Figure 2E).

**Figure 2.**
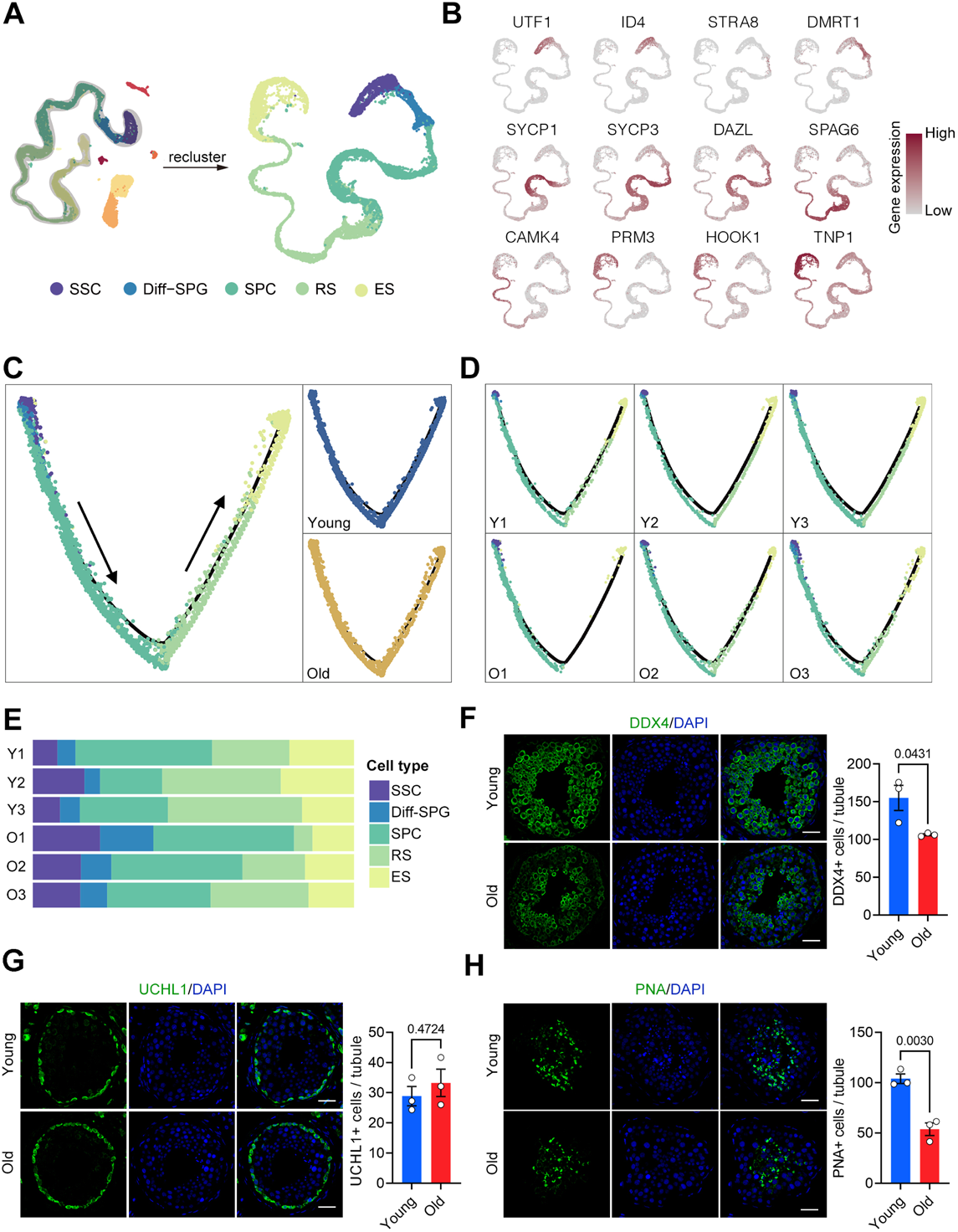
Aging-related Cellular Alterations Along the Trajectories of Spermatogenesis. (A) UMAP projection of germ cells reveals the developmental progression of spermatogenesis in young and old human testis. (B) Expression patterns of known spermatogenic markers projected onto the UMAP plot. The color key, ranging from gray to brown, indicates low to high gene expression levels, respectively. (C) Pseudotime trajectory (Monocle analysis) of germ cells. Cells are colored according to the cell type and group. Arrows indicate the developmental stages from SSCs to ESs. (D) Deconvolution of the Monocle pseudotime plot according to the donors of origin. (E) Relative proportions of cells at different spermatogenic stages in the samples analyzed. (F) Immunostaining and quantification of DDX4 in young and old human testis sections. Left, representative image of DDX4 in human testis sections. Right, quantification of the number of DDX4-positive cells per seminiferous tubule. Scale bars, 35 μm. Young, n=3; Old, n=3. Data are expressed as mean ± sem. Significance was determined by Student’s t-test. (G) Immunostaining and quantification of UCHL1 in young and old human testis sections. Left, representative image of UCHL1 in human testis sections. Right, quantification of UCHL1+ cells per seminiferous tubule. Scale bars, 35 μm. Young, n=3; Old, n=3. Data are expressed as mean ± sem. Significance was determined by Student’s t-test. (H) Immunostaining and quantification of PNA in young and old human testis sections. Left, representative image of PNA in human testis sections. Right, quantification of PNA+ cells per seminiferous tubule. Scale bars, 35 μm. Young, n=3; Old, n=3. Data are expressed as mean ± sem. Significance was determined by Student’s t-test.

Immunofluorescence analysis of the germ cell marker, DEAD-box helicase 4 (DDX4), supported the notion that germ cells were reduced in the testes of the old group (Figure 2F). Further analysis revealed that the numbers of UCHL1+ SSCs were equivalent between the young and old groups (Figure 2G), indicating that a change in the quantity of SSCs might not be largely involved in the aging-associated deficiency of spermatogenesis. The number of peanut agglutinin (PNA)+ RSs and ESs was significantly lower in testes of young versus elderly individuals (Figure 2H). This indicated that germ cell differentiation was severely impacted by aging, which is consistent with our histological examinations (Figure 1A).

Taken together, our transcriptomic data combined with the results of our immunofluorescent studies indicate that there are cellular differences in spermatogenesis-related cells of the young and old groups, and that RSs and ESs decreased considerably with age.

### Aging-related molecular alterations along the trajectories of spermatogenesis

We next focused on age-related transcriptional changes in the identified germline cell types. Aging has been associated with increased transcriptional noise(Nikopoulou, Parekh, & Tessarz, 2019). Here, calculation of the age-relevant coefficient of variation (CV) revealed that SPCs, Diff-SPGs, and SSCs exhibited higher transcriptional noise than later-stage germline cells (Figure 3A), indicating that aging caused higher variability in early-stage germline cells compared to late-stage germline cells. We further identified 174, 102, 90, 644, and 214 upregulated genes and 112, 46, 127, 891, and 665 downregulated genes (|avg_logFC| > 0.25 and p-value < 0.05) in the SSC, Diff-SPG, SPCs, RS, and ES germline subtypes, respectively, in the old versus young groups (Figure 4B; Supplementary file 2). Notably, only ∼22% of the differentially expressed genes (DEGs) were shared by at least two cell populations; the majority of DEGs were cell-type-specific, indicating that aging has stage-specific effects in this setting. We therefore performed GO analysis based on gene set enrichment analysis (GSEA), with the goal of exploring the aging-associated alterations in the cellular functions of each germ cell type (Figure 3C, S3A-S3D; Supplementary file 3). Given that SSCs represent the only germline stem cells and play essential roles in long-term spermatogenesis(Di Persio et al., 2021), we focused on the aging-associated changes of gene expression in SSCs. We observed that the genes downregulated in SSCs of the old group were associated with base-excision repair (BER) and nucleotide-excision repair (NER), two important excision repair mechanisms for DNA damage(Ray Chaudhuri & Nussenzweig, 2017) (Figure 3C). Therefore, we performed gene set-score analysis for the BER and NER signaling pathway of SSCs in the young and old groups (Supplementary file 4). We observed a prominent decrease of the gene-set score of the BER pathway in aged SSCs, compared to young SSCs, whereas that of the NER pathway did not significantly differ between the two groups (Figure 3D). Moreover, we found that a number of BER-promoting genes, including NTHL1, NEIL2, MPG, APEX1, PARP1, and POLD2, were transcriptionally downregulated in SSCs from old testes (Figure 3E). Consistent with these detected changes in mRNA levels, immunostaining analyses confirmed that the protein levels of NTHL1 and APEX1 were decreased in SSCs of the old group compared to the young group (Figures 3F, 3G), further supporting the idea that the DNA repair function of SSCs is compromised during human testicular aging.

**Figure 3.**
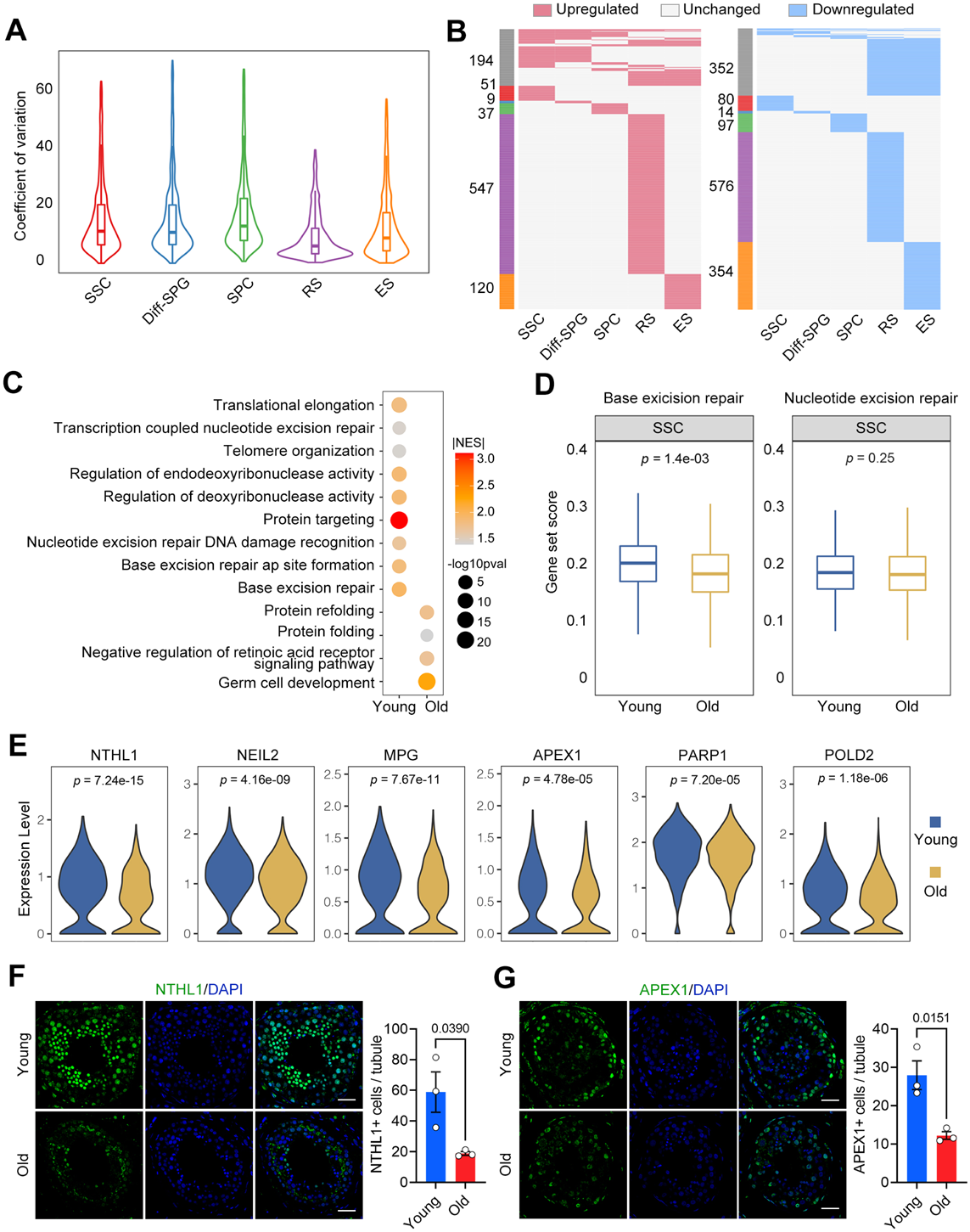
Aging-related Molecular Alterations Along the Trajectories of the Spermatogenesis. (A) CV analysis showing the aging-associated transcriptional noise in germ cells. (B) Heatmaps showing the distribution of upregulated (red) and downregulated (blue) DEGs (log2FC>0.25, min.diff.pct = 0.1, *p*-value <0.05) between old and young human germ cells. Genes that are not differentially expressed are indicated in gray, and the numbers of DEGs are indicated. The gray bars on the left of the heatmaps denote DEGs shared by at least two cell types, and the others are cell-specific DEGs. (C) Representative GO terms of DEGs in old and young human SSCs. Dot size indicates the range of *p*-value. Color keys, ranging from gray to orange to red, indicate the absolute value of the normalized enrichment score (NES). (D) Gene set score analysis for the BER and NER signaling pathways of SSCs from the young and old groups. Wilcoxon rank sum test was used; *p*-value is indicated. (E) Violin plot showing expression levels of BER-associated genes for SSCs in young and old samples. (F) Immunostaining and quantification of NTHL1 in young and old human testis sections. Left, representative image of NTHL1 in human testis sections. Right, quantification of NTHL1-positive cells in seminiferous tubules of young and old human testis sections. Scale bars, 35 μm. Young, n=3; Old, n=3. Data are expressed as mean ± sem. Significance was determined by Student’s t-test. (G) Immunostaining and quantification of APEX1 in young and old human testis sections. Left, representative image of APEX1 in human testis sections. Right, quantification of APEX1+ cells in seminiferous tubules of young and old human testis sections. Scale bars, 35 μm. Young, n=3; Old, n=3. Data are expressed as mean ± sem. Significance was determined by Student’s t-test.

**Figure 4.**
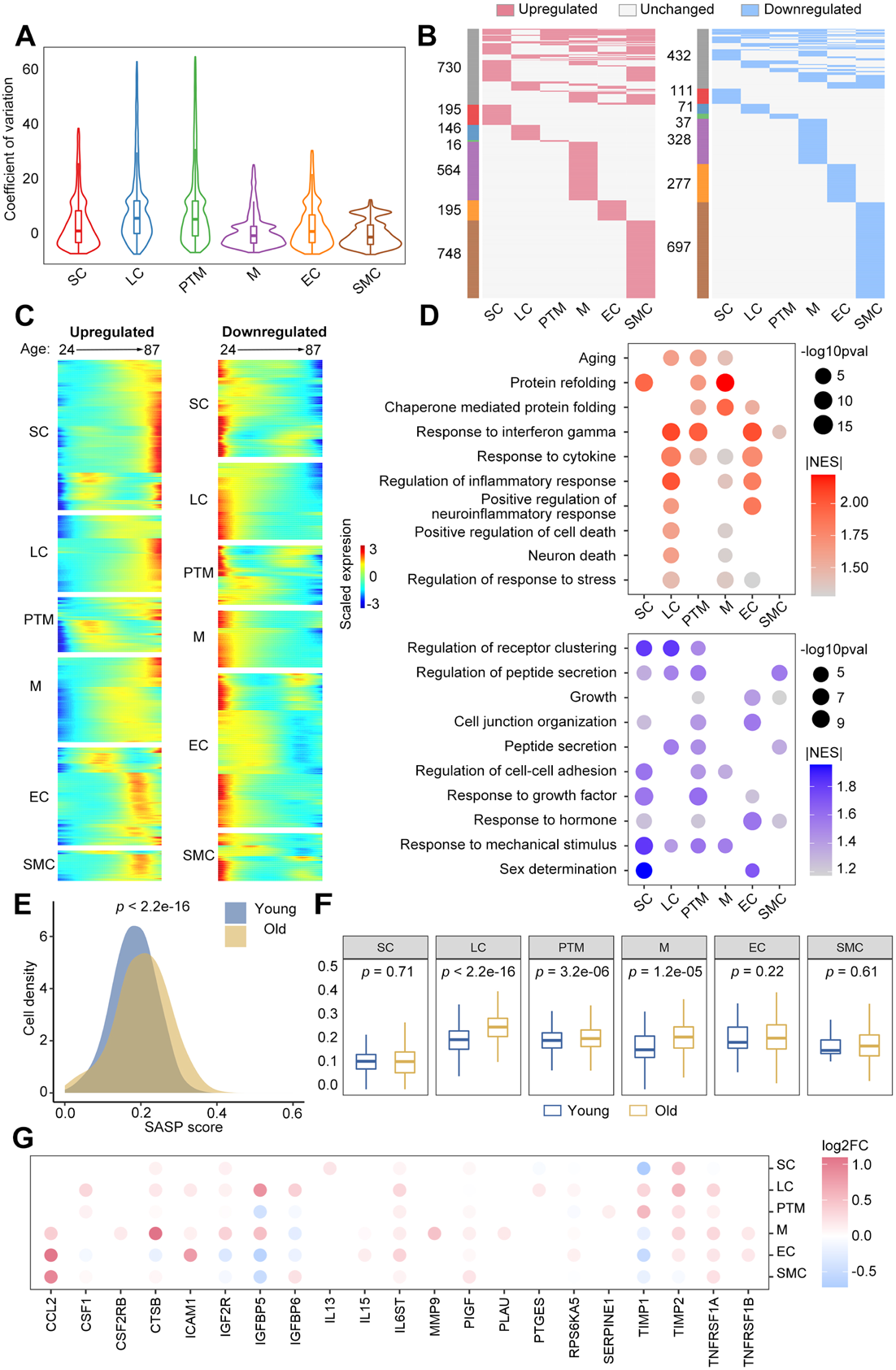
Changes in the Transcriptional Profiles of Somatic Cells during Human Testicular Aging. (A) CV analysis showing the aging-associated transcriptional noise in somatic cells. (B) Heatmaps showing the distribution of upregulated (red) and downregulated (blue) DEGs (|log2FC|>0.25, min.diff.pct = 0.1, *p*-value <0.05) between old and young human germ cells. Genes not differentially expressed are in gray, and the numbers of DEGs are indicated. (C) Heatmaps showing the age-related upregulated (left) and downregulated (right) DEGs of somatic cell types. The color key, ranging from blue to red, indicates low to high gene expression levels. (D) Representative shared GO terms of age-related upregulated (top) and downregulated (bottom) DEGs in different somatic cell types. Dot size indicates the range of *p*-value. The color keys, ranging from gray to red (top) or from gray to blue (bottom), indicate the absolute values of the normalized enrichment score (NES). (E) Density plot showing the distribution of cells with different SASP scores in the young and old groups. Wilcoxon rank sum test was used; *p*-value is indicated. (F) Box plot showing SASP scores in different types of somatic cells. Wilcoxon rank sum test was used; *p*- values are indicated. (G) Dot plot showing the log2-transformed fold change of SASP-related genes in somatic cells from young versus old groups. Only genes showing a statistically significant difference between the young and old groups are shown. Dot color indicates the log2-transformed fold change.

As BER is a major pathway for repairing oxidative base damage, alkylation damage, and abasic sites on DNA(Ray Chaudhuri & Nussenzweig, 2017), its dysregulation in SSCs is likely to contribute to the age-related increase of de novo germline mutations in males.

### Changes in the transcriptional profiles of somatic cells during human testicular aging

We next quantified the populations of the main somatic cell types, including LCs, SCs, and PTMs, in young and old testes by immunofluorescence analysis (Figure S4A-C). To further explore the mechanisms of testicular aging at the cellular level, we compared the gene expression patterns in somatic cells between the groups. By calculating the age-relevant CV(S. Wang et al., 2020), we found that the aging-accumulated transcriptional noise was higher in LCs, PTMs, and SCs compared to the other somatic cell types (Figure 4A). This suggested that LCs, PTMs, and SCs may be more vulnerable to age-related stress than the other somatic cell types. We next explored aging-associated DEGs in somatic cells between the groups. We identified thousands of genes that represented DEGs (|avg_logFC| > 0.25 and *p*-value < 0.05) in at least one somatic cell type of human testis during aging (Figures 4B). Further analysis enabled us to identify 621, 316, 228, 944, 447, and 1274 upregulated genes and 280, 225, 171, 567, 447, and 913 downregulated genes in SCs, LCs, PTMs, Ms, ECs, and SMCs, respectively, in the old versus young comparison (Figure 4B; Supplementary file 2. Notably, only ∼26% of the DEGs were shared by at least two cell populations, indicating that the effects of aging are largely somatic cell type-specific in this setting.

To identify DEGs that constantly increased or decreased in somatic cells during aging, we aligned datasets of individual samples by their chronological age and explored the expression patterns of 2594 DEGs that were upregulated with age and 1953 DEGs that were downregulated with age in six somatic cell types (Figure 4C). GO enrichment analysis revealed that the genes upregulated with age were mainly associated with cytokine and stress responses, unfolded proteins, and apoptotic signaling. In comparison, the genes downregulated with age were mainly related to peptide secretion, hormone responses, and growth (Figure 4D; Supplementary file 3). As the senescence-associated secretory phenotype (SASP) is a common feature of senescent cells and usually contributes to a low-grade inflammatory state(Baker & Petersen, 2018), we questioned whether the somatic niche in the aged testis might present an elevated SASP environment. Indeed, our analysis revealed that somatic cells exhibited much higher gene-set scores for SASP (Figure 4E). Notably, LCs, PTMs and Ms showed elevated SASP scores with age (Figure 4F, 4G; Supplementary file 4), indicating that these three cell types had strong contributions to the age-related inflammatory state in human testis.

Next, we performed an integrative comparative analysis of aging-associated DEGs with aging/longevity-associated genes from the GenAge database(Tacutu et al., 2018) and further found that many aging/longevity-associated genes were differentially expressed in one or more somatic cell type (Figure S4D). For instance, the molecular marker of senescent cells, CDKN1A, was upregulated in LCs, PTMs, and ECs, whereas several cell survival-related genes (PLCG2, IGFBP2, and GSTP1) were downregulated in three or more somatic cell populations during testicular aging (Figures S4D).

Collectively, these results indicate that a series of aging-related molecular changes occur in aged human testicular somatic cells and likely contribute to testicular homeostasis and aging.

### Dysfunction of LCs during human testicular aging

LCs produce testosterone and are thus critical for reproductive function and general health(Zirkin & Papadopoulos, 2018). LC dysfunction causes testosterone deficiency, arrested spermatogenesis, and infertility. Here, we observed the highest level of aging-accumulated transcriptional noise in LCs (Figure 4A). Thus, we next focused on the aging-associated changes of gene expression in LCs. For functional validation, we assessed the ability of aged LCs to produce testosterone. We isolated primary LCs from human testicular tissues of both groups (Figure S5A, B). Assessment of testosterone levels in the media revealed that LCs from the young group produced significantly more testosterone than those from the old group (Figure 5A). Consistently, when we cultured small pieces of testicular tissue in vitro, testosterone production was significantly lower in the old group compared with the young group (Figure 5B). These results demonstrate that the basic function of LCs appears to become compromised during aging.

**Figure 5.**
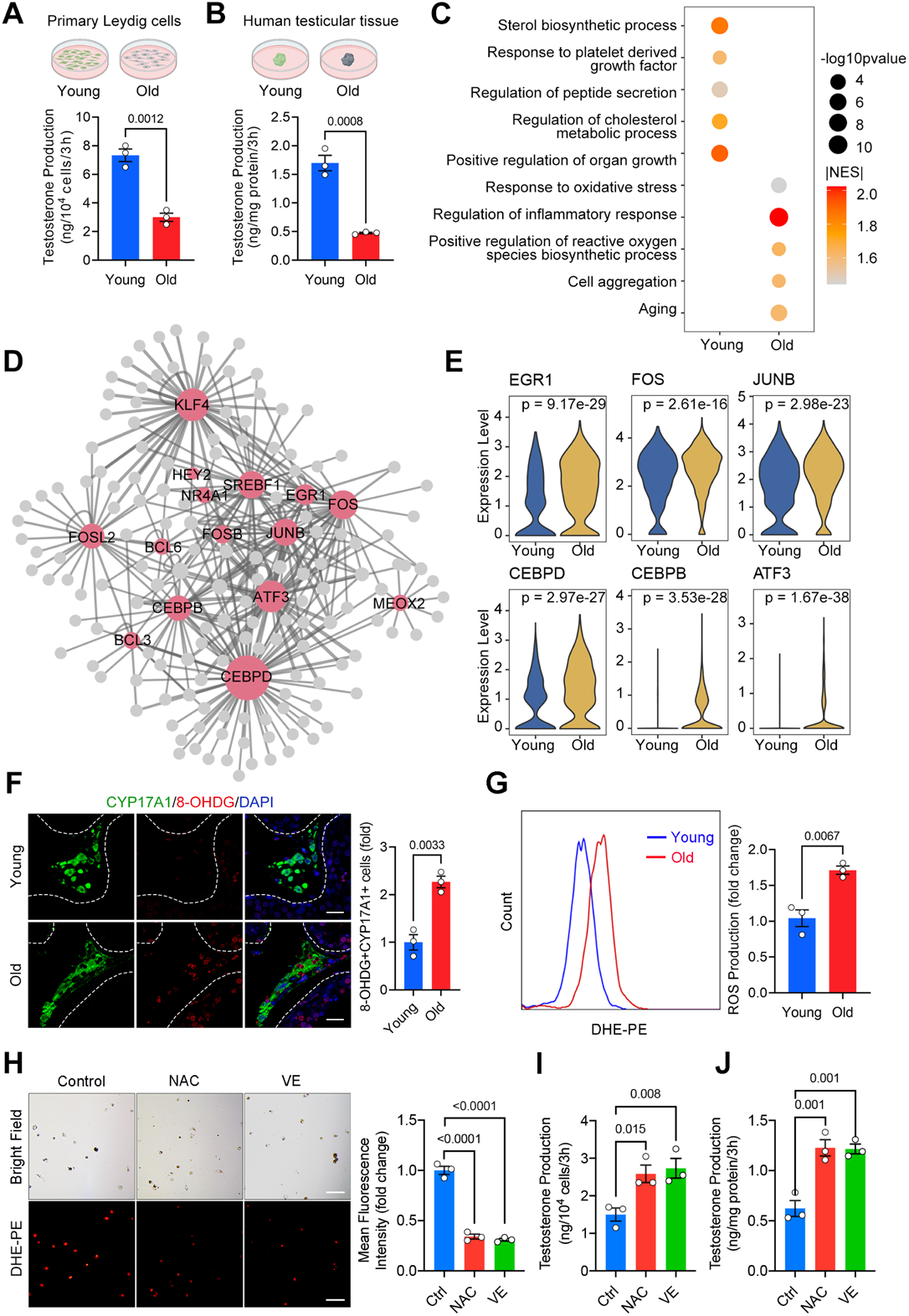
Dysfunction of Leydig Cells During Human Testicular Aging. (A,B) Quantification of testosterone production in primary LCs (A) and human testicular tissue (B). Young, n=3; Old, n=3. Data are expressed as mean ± sem. Significance was determined by Student’s t-test. (C) Representative GO terms of DEGs in old and young human LCs. Dot size indicates the range of *p*-value. Color keys, ranging from gray to orange to red, indicate the absolute values of the normalized enrichment score (NES). (D) Network visualization of potential key transcriptional regulators of upregulated DEGs in LCs. Node size is positively correlated with the number of directed edges. Edge width is positively correlated with the NES score. Transcription factors are highlighted in red; others are in gray. (E) Violin plot showing the expression levels of age-associated upregulated transcriptional factors. Wilcoxon rank sum test was used; *p*-value is indicated. (F) Immunostaining and quantification of 8-OHDG and CYP17A1 in young and old human testis sections. Left, representative image of 8-OHDG and CYP17A1 in human testis sections. Right, quantification of the proportion of 8-OHDG+ LCs (CYP17A1+). Scale bars, 25 μm. Young, n=3; Old, n=3. Data are expressed as mean ± sem. Significance was determined by Student’s t-test. (G) Quantification of ROS production in young and old primary LCs, as measured by flow cytometry. Young, n=3; Old, n=3. Data are expressed as mean ± sem. Significance was determined by Student’s t-test. \*\**p<*0.01. (H) Representative images and quantification ROS production by primary human LCs isolated from old testes, comparing control with antioxidant-treated (NAC or VE) groups. Left, representative bright field and immunofluorescent (DHE-PE) images of primary LCs. Right, quantification of ROS production by primary LCs. Scale bars, 100 μm. Control, n=3; NAC (10 mM), n=3; VE (5 μM), n=3. Data are expressed as mean ± sem. Significance was determined by one-way ANOVA. (I) Quantification of testosterone production by primary human LCs isolated from old testes in control and antioxidants treatment (NAC and VE) groups. Control, n=3; NAC (10 mM), n=3; VE (5 μM), n=3. Data are expressed as mean ± sem. Significance was determined by one-way ANOVA. \**p*<0.05, \*\**p*<0.01. (J) Quantification of testosterone production by old human testicular tissues in control and antioxidant-treated (NAC and VE) groups. Control, n=3; NAC (10 mM), n=3; VE (5 μM), n=3. Data are expressed as mean ± sem. Significance was determined by one-way ANOVA.

To further characterize the changes in LCs during human testicular aging, we performed GO enrichment analysis based on GSEA analysis. Our results revealed that the genes upregulated with age were mainly associated with regulating the response to reactive oxygen species (ROS) and cell aggregation (Figure 5C). Notably, genes related to inflammatory responses and aging were also upregulated in aged human LCs (Figure 5C), which is consistent with their elevated SASP score and expression of the senescence marker, CDKN1A (Figure 4F, S4D). By comparison, the downregulated DEGs were mainly enriched in GO terms such as cholesterol metabolic processes and the response to platelet-derived growth factor; these findings are consistent with the tendencies of aged LCs to exhibit decreases in testosterone production and cell number, respectively (Figure 5A-C; S4A). Genes related to positive regulation of organ growth (IGF1, IGF2, ARX, DDX39B, AKAP6, TBX2, and ZFPM2) were also downregulated in aged human LCs, likely contributing to testicular aging.

To identify critical regulators linked to LC aging, we constructed gene-regulatory networks based on aging-associated DEGs. We noticed that several TFs, such as CEBPD, FOSL2, JUNB, KLF4, ATF3, and EGR1, were among the top upregulated genes in aged human LCs (Figure 5D; Supplementary file 5). This finding was in agreement with the prominent upregulation of the response to oxidative stress found in our GO analysis, as EGR1 is a well-known central regulator of the oxidative stress response(Dai et al., 2021). Additional oxidative stress regulators, such as CEBPD(S. M. Wang et al., 2018), ATF3(Feng, Li, Jin, Gong, & Xia, 2021) (Figure 5E), and JUNB(M. Chen, Li, Shi, Zhang, & Xu, 2019), were also upregulated (Figure 5E), further supporting the notion that redox homeostasis is impaired in aged human LCs.

Consistent with the transcriptomic changes, we observed an increase in cells positive for 8-OHDG, which is a recognized biomarker of oxidative stress(S. Wang et al., 2020; Zou et al., 2021), in aged LCs compared with young LCs (Figures 5F). Moreover, primary human LCs from the old group generated more intracellular ROS than those from the young group (Figures 5G). Of note, 8-OHDG-positive cells also detected in several other testicular cell types (Figures 5F), suggesting that oxidative stress is involved in the aging process of human testis.

Collectively, our data suggest that age-related impairment in redox homeostasis may function as a molecular basis for LC dysfunction, and thereby likely contributes to late-onset hypogonadism during male aging.

### Antioxidants restore the testosterone production of LCs from aging human testis

Next, we asked whether the suppression of oxidative stress could counteract the cellular dysfunction of LCs. As small-molecule agents, such as N-acetylcysteine (NAC) and vitamin E (VE), have been reported to reduce oxidants(H. Chen et al., 2005; W. Zhao et al., 2017), we tested the effect of these antioxidants on the function of primary LCs and testicular samples from the old group. Not surprisingly, primary LCs treated with NAC or VE exhibited decreases in the ROS level (Figures 5H, S5C). Interestingly, however, the antioxidant treatments recovered testosterone production in isolated primary LCs, whereas the vehicle control did not (Figure 5I). The antioxidant treatments also restored testosterone production in aged human testicular samples (Figure 5J), suggesting that antioxidative strategies may rejuvenate LCs and recover testosterone production in elderly humans. Collectively, these findings suggest a new platform for uncovering potential intervention targets and compounds for alleviating late-onset hypogonadism and testicular aging.

### Cell-cell communication changes during human testicular aging

The maintenance of multicellular organism morphology and function largely relies on cell-cell communication. Therefore, using the ligand–receptor interaction tool CellChat(Jin et al., 2021), we next systematically inferred changes in communication networks between testicular cells at the single-cell level during aging. To our surprise, the interaction number and interaction strength were enhanced in the old group compared to the young group (Figure 6A; Supplementary file 6). We then assessed differential interactions of testicular cells as either signal sources or signal targets during aging. LCs and PTMs showed similar and relative decreases in their communicating roles during aging; meanwhile, there was a global trend for increasing interactions in old testis compared to young testis (Figure 6B). Consistently, investigation of the outgoing and incoming signaling patterns revealed that there were relative declines of the outgoing interaction strengths for aged LCs and PTMs compared to their young counterparts (Figure 6C). These results encouraged us to further study the potential role of LCs and PTMs in testicular aging.

**Figure 6.**
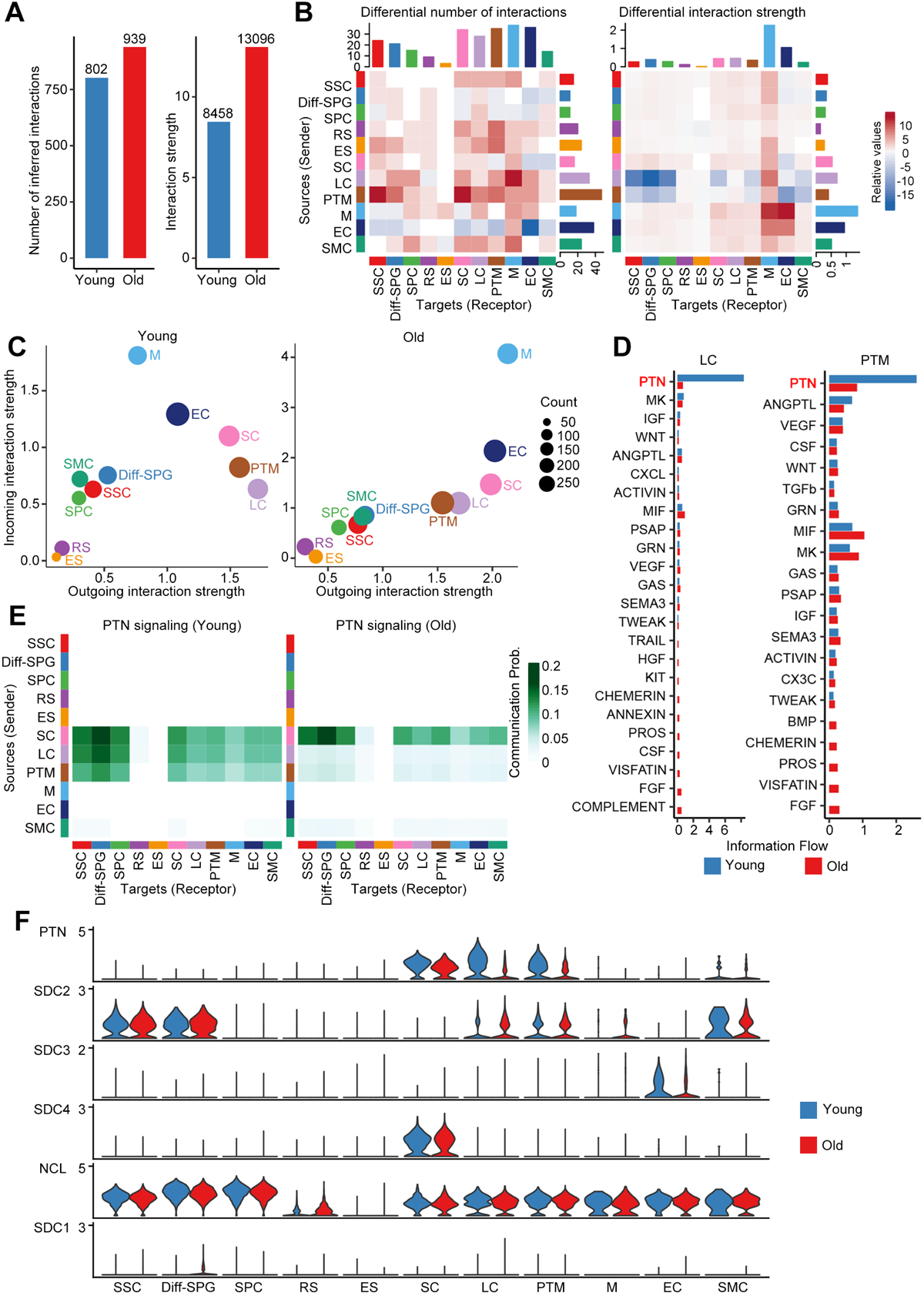
Cell-cell communication changes during human testicular aging. (A) Bar plot showing the number (left) or strength (right) of interactions in the cell-cell communication network, as analyzed by CellChat. (B) Heatmap showing the different number (left) or strength (right) of interactions in the cell-cell communication network between young and old groups. Red or blue colors represent increased or decreased signaling, respectively, in old groups compared to young. (C) The outgoing (sending signals) and incoming (receiving signals) interaction strength of the cell populations in young (left) and old (right) groups. “Count” indicates the number of inferred links (both outgoing and incoming) associated with each cell group. (D) Information flow of significant signaling pathways sent by LCs (left) or PTMs (right) and received by all cell populations in young (blue columns) and old (red columns) groups. The top signaling pathway, PTN, is labeled in red. (E) Heatmap showing the inferred intercellular communication networks of the PTN signaling pathways in young (left) and old (right) cell populations. Color key, ranging from white to green, indicates the between-cell communication probability for the PTN signaling pathway. (F) Violin plot showing expression levels of ligand-receptor pairs for the PTN signaling network in young and old cell populations.

When we compared the information flow for each signaling pathway sent by LCs or PTMs between the young and old groups, we found that the pleiotrophin (PTN) signaling pathway ranked highest in flow for young testis and declined prominently during aging (Figure 6D). Based on this finding, we investigated the PTN signaling pathway network of testicular cells. In the young group, PTN signaling was mainly sent by SCs, LCs, and PTMs and received by SSCs, Diff-SPGs, SPCs, and somatic cells. In the old group, in comparison, PTN signaling was dramatically lower between LCs, PTMs and other cells, but relatively similar between SCs and other cells (Figure 6E).

In the testis, growth factors and morphogen signals play important roles in spermatogenesis. We next inferred the age-related changes in the communication networks of target germ cells. Similar to the above-described findings, PTN was prominent among the incoming signal pathways of SSCs, Diff-SPGs, and SPCs (Figure S6A). This supports the idea that PTN signaling forms a potential regulatory axis between somatic cells and spermatogenesis. Our analysis identified five significant ligand-receptor pairs for PTN signaling: PTN-NCL, PTN-SDC1, PTN-SDC2, PTN-SDC3, and PTN-SDC4. Among them, PTN-NCL contributed most highly to PTN signaling in both young and old groups (Figure S6B, S6C). The expression level of PTN was downregulated in LCs and PTMs of the old group compared to the young group, whereas the expression levels of NCL, SDC1, SDC2, SDC3, and SDC4 did not differ with age in most of the other tested cell populations (Figure 6F). This further supports the importance of LCs and PTMs in human testicular aging. Notably, PTN was previously reported to maintain stemness and activate cell proliferation, and to play important roles in neural development, angiogenesis, and bone development(Deuel, Zhang, Yeh, Silos-Santiago, & Wang, 2002). The downregulation of PTN and PTN pathway signaling during aging may help explain age-related male fertility declines and hypogonadism.

## Discussion

In this report, we present the single-cell survey of human testicular aging, providing insights into the mechanisms by which human testes age. Our analyses provide four noteworthy contributions. First, we elucidated the gene expression signatures of 11 types of human testicular cells (including germline and somatic cells) and identified several previously unreported cell type-specific markers. Second, analysis of age-associated gene expression changes revealed that DNA repair genes are downregulated in aged human SSCs, which may represent a driver for the age-related increase of de novo germline mutations in males. Third, human LCs exhibited aging-associated upregulation of oxidative stress genes, and antioxidant treatment recovered the functions of aged human LCs. Fourth, our results suggested that decreases in PTN signaling contribute to testicular aging. Together, these observations provide novel insights into human testicular aging and identify new potential targets for treating human disorders associated with testicular aging.

Although the fertility decline and testosterone deficiency caused by testicular aging undermine reproductive and general health in aging males, the impact of age on the human testis is still poorly understood(Kaufman et al., 2019; Santiago et al., 2019). The previous studies have mainly detailed the morphological changes that occur in the human testis during aging, while the molecular mechanisms underlying testicular aging have remained largely unknown(Jiang et al., 2014; Mularoni et al., 2020; Perheentupa & Huhtaniemi, 2009). Moreover, the few previous mechanistic reports have used analytical methods with inherent biases that limit their ability to provide information for all cell populations(Han, Hong, Lee, Hong, & Cho, 2021; Stockl et al., 2021). To address these gaps, we herein used scRNA-seq analysis, which enables the study of heterogeneous cells and the identification of cell states and cell type-specific gene changes during aging or disease emergence(Di Persio et al., 2021; Guo et al., 2021; S. Wang et al., 2020; Yang et al., 2022). Recent scRNA-seq-based studies have reported transcriptomic and functional changes in immune cells during aging(Zheng et al., 2020) and in tissues such as the central nervous system(Ximerakis et al., 2019), mammary gland(C. M. Li et al., 2020), and ovary(S. Wang et al., 2020). scRNA-seq was also previously used to analyze human testes, and the results improved our understanding of testicular biology and related pathology(Alfano et al., 2021; Di Persio et al., 2021; Guo et al., 2018; Guo et al., 2020; Guo et al., 2021). For instance, scRNA-seq analyses of testes revealed profiles for human testis development(Guo et al., 2020; Guo et al., 2021), Klinefelter syndrome(Mahyari et al., 2021), idiopathic germ cell aplasia(Alfano et al., 2021), and azoospermia(L. Zhao et al., 2020). Using the approach, a recent study of human testicular aging has been reported, providing candidate molecular mechanisms of the testicular changes during aging(Nie et al., 2022). However, the critical molecular drivers underlying germ cell and Leydig cell functional decline during aging remain unclear. Here, we used scRNA-seq to comprehensively delineate cell type-specific, age-associated gene expression changes in human testes. To our knowledge, this is the first scRNA-seq study to reveal the cell type particularly vulnerable to aging and uncover the potential molecular drivers of testicular aging by in-depth sequencing and reproducible bioinformatics tools.

Since human male germ cells have a unique germline-specific aging process(Laurentino et al., 2020; Pohl et al., 2019), we first analyzed germline cells and identified five cell categories based on their specific scRNA-seq signatures. We show that human germline aging is linked to decreased RSs and ESs in aged testes, consistent with previous reports that aging males showed a decline in sperm parameters(Pohl, Gromoll, Wistuba, & Laurentino, 2021; Pohl et al., 2019). SSCs are known to be the foundational unit of fertility in all male mammals(Sharma et al., 2019; Tan & Wilkinson, 2020), but the research has been slow to advance our knowledge of human SSC biology, especially in the aging process. Here, we characterized the molecular changes in human SSCs during testicular aging. Surprisingly, we found that BER, which is a major pathway for DNA repair(Ray Chaudhuri & Nussenzweig, 2017), was downregulated in aged human SSCs. Consistently, immunostaining revealed age-related decreases in NTLH1 and APEX1, which are important BER(Galick et al., 2013; M. Li et al., 2018), in human SSC populations. Cells are exposed to various endogenous and exogenous insults that induce DNA damage that, if left unrepaired, can impair genomic integrity and lead to the development of various diseases(Ray Chaudhuri & Nussenzweig, 2017). Among the germline cells, differentiating germ cells exist for only the duration of one spermatogenic cycle, but SSCs are long-living stem cells that exist throughout the male’s life. Thus, SSCs are susceptible to age- accumulated DNA damage(Laurentino et al., 2020). Indeed, whole-genome sequencing studies have shown that de novo point mutations in children arise predominantly from the male SSCs, and the mutational frequency increases with paternal age(Maher et al., 2016). These mutations are linked to increased risks of breast cancer, developmental disorders, behavioral disorders, and neurological disease in the children of older men(Maher et al., 2016; Yatsenko & Turek, 2018). These findings support the hypothesis that disturbance of BER in SSCs drives the age- related increase of de novo germline mutations and may have further negative consequences for the offspring’s health. These possibilities remain to be thoroughly assessed in future studies. Based on the derived dataset, we identified thousands of cell type-specific DEGs that highlighted the molecular changes underlying the aging of human testicular somatic cells. Specifically, the upregulation of stress responses and apoptotic signaling and the downregulation of growth-related pathways were prominent features of somatic cells in aged testes. Moreover, somatic cells exhibited much higher SASP-related gene expression levels, suggesting that the SSC niche may be destroyed with age. LCs produce a large amount of ROS when testosterone is synthesized, making them vulnerable to ROS-induced damage(Cao, Leers- Sucheta, & Azhar, 2004; Zirkin & Papadopoulos, 2018). Accordingly, we found that the aging- associated DEGs were enriched for the ROS response and apoptosis pathways in LCs. Intriguingly, we also observed an elevated inflammatory response in aged LCs, which may reflect an accumulation of ROS-induced macromolecule damage(Forman & Zhang, 2021). Given that inflammation is an adaptive response to noxious stress or malfunction, the elevated inflammation we observed in aged LCs implies that intracellular homeostasis becomes skewed toward a chronic stress state with age. Recent evidence suggests that oxidative stress may be linked to LC dysfunction and hypogonadism in rodents(Cao et al., 2004; H. Chen et al., 2005). Consistently, we found that DNA oxidation markers were increased in aged LCs compared to young LCs. Moreover, isolated primary human LCs from the old group generated more intracellular ROS than LCs from young group, supporting the notion that oxidative stress induces LC dysfunction in aging human testes.

Based on our present findings, we speculated that oxidative stress could be targeted as a therapeutic avenue to prevent LC dysfunction in aging males. A number of reports have examined the beneficial effects of antioxidants on the function of murine LCs, but mostly under pathological conditions (e.g., testicular torsion or diabetes) or following exposure to various toxic agents(H. Chen et al., 2005). To our knowledge, no previous study has reported an antioxidant intervention that successfully improved human LC function. Here, we show that antioxidants restored testosterone production not only in primary human LCs but also in human testicular samples from the old group, suggesting that scRNA-seq could represent a new platform for uncovering intervention targets and compounds for alleviating late-onset hypogonadism and testicular aging in humans.

Testicular physiology broadly relies on cell-cell signaling, and imbalances in this signaling most likely contribute to various forms of spermatogenic impairment(Di Persio et al., 2021; Mahyari et al., 2021). Here, we found that LCs and PTMs showed decreases in their communication roles during aging, which contrasted with the global tendency toward increased interaction in old testis relative to young testis. This suggested that LCs and PTMs play important roles in human testicular aging. Moreover, we found that the PTN signaling pathway was top-ranked in young testis and declined dramatically in importance among LCs, PTMs, and other cells of the old group. PTN is an 18-kDa heparin-binding secretory growth/differentiation factor for different cell types that was reported to be expressed at only low levels in some cells of the brain, bones, gut, uterus, and ovary(Deuel et al., 2002). However, PTN was found to be expressed at a significantly higher level in cells of the testis, especially LCs(Vanderwinden, Mailleux, Schiffmann, & Vanderhaeghen, 1992). A previous study showed that a dominant-negative PTN mutant caused testicular atrophy and apoptosis among spermatocytes at all stages of development; this suggested that PTN plays a central role in normal spermatogenesis, and that interruption of PTN signaling may lead to sterility in males(Zhang, Yeh, Zhong, Li, & Deuel, 1999). It is possible that the continuous self-renewal and differentiation of developing spermatogonia make them uniquely susceptible to the loss of PTN signaling. However, the effects of PTN on human testicular aging are not yet fully understood and need to be further elucidated in future studies.

In summary, the present study provides the comprehensive single-cell transcriptomic atlas of young and aged human testis and broadens our understanding of cell type-specific gene signatures in the human testis. Importantly, our work offers insights into the molecular mechanisms underlying testicular aging in humans, which could help the field work toward developing targeted interventions to protect against testicular aging and/or suggest new tools to rejuvenate aged germline and Leydig cells.

## Materials and methods

### Human testicular tissues

Human testis samples were obtained from 6 male donors who underwent testicular excision or biopsy for the following indications: testicular teratoma (Y1; 28 years old), testicular Leydig cell tumor (Y2; 24 years old), obstructive azoospermia (Y3; 31 years old), testicular cyst (O1; 61 years old), and prostate cancer without androgen deprivation therapy (O2; 87 years old; O3; 70 years old). For patients Y1, Y2, and O1, we collected normal tissues distant from lesions. The old donors were confirmed to have offspring, which was taken as indicating that they had normal reproductive function when young. Informed consent was obtained from all of the above-listed patients.

### Tissue processing

After being collected from the operating room, the samples were transported to the laboratory on ice in storage solution (Miltenyi Biotec, Shanghai, China) within 1 h. The tunica was removed and testicular tissues were minced and washed three times with phosphate-buffered saline (PBS) to eliminate the storage solution and blood. To ensure accuracy and stability, ∼200 mg of tissue was immediately applied for scRNA-seq. Thereafter, tissue samples (∼500 mg) were fixed with 4% paraformaldehyde (PFA; Thermo Fisher Scientific, Wilmington, DE, USA) for histochemistry or immunostaining analyses. The remaining testis tissues were cryopreserved for functional assessments, as previously described(Guo et al., 2018).

### Sample preparation for scRNA-seq

For single-cell sequencing, testicular samples were minced and subjected to a standard two-step digestion procedure(Guo et al., 2018). Firstly, the tissues were digested with dissociation buffer including 1 mg/mL type IV collagenase (Gibco, Grand Island, NY, USA) and 200 μg/mL DNase I (Roche, Indianapolis, IN, USA) dissolved in DMEM/F-12 (Gibco) at 37℃ in a water bath for 15 min. The tissues were gently pipetted against the bottom of tube with a Pasteur pipet, passed through a 40-μm filter, washed twice with PBS, and stored temporarily at 4℃. Given that this strategy might not have isolated all cells present in the spermatogenic tubule, we redigested the samples left on the filter with 0.25% Trypsin-EDTA at 37℃ in a water bath for 10 min. The digestion was terminated with termination buffer containing 10% fetal bovine serum (FBS; Thermo Fisher Scientific, Wilmington, DE, USA) and the samples were filtered through a 40-μm filter and centrifuged. Finally, the obtained cells were combined, passed through a 40-μm filter, and resuspended in PBS. Cell numbers were counted with a Cellometer Auto T4 automated cell counter (Nexcelom Bioscience, Lawrence, MA, USA) and resuspended at 1000 cells/μL in PBS containing 0.1% BSA for single-cell sequencing.

### H&E staining

Fixed tissues were embedded in paraffin and sectioned at 4 μm. The sections were deparaffinized with xylene, rehydrated with an ethanol series (100%, 95%, 85%, 75%), and stained with hematoxylin and eosin. Images were collected with a DMi8 microscope (Leica, Wetzlar, Germany).

### Masson staining

Fixed tissues were embedded in paraffin and sectioned at 4 μm. After being deparaffinized with xylene and rehydrated with an ethanol series (100%, 95%, 85%, 75%), the sections were stained overnight with potassium bichromate solution and rinsed with running tap water for 5 min. The sections were then incubated sequentially in Weigert’s iron hematoxylin working solution for 10 min, Biebrich scarlet-acid fuchsin solution for 10 min, and phosphomolybdic-phosphotungstic acid solution for 15 min. Between each step, the slides were washed in distilled water. The sections were then transferred directly (without being rinsed) to aniline blue solution and stained for 10 min. After being briefly washed with distilled water, the sections were differentiated in 1% acetic acid solution for 5 min, and then rinsed with distilled water. Finally, the sections were dehydrated and mounted using resinous mounting medium. Images were collected with a DMi8 microscope (Leica).

### Single-cell RNA-seq library construction, sequencing, and alignment

Single-cell suspensions were loaded to a 10x Chromium Controller instrument (10× Genomics, Pleasanton, CA, USA) and ∼5000 single cells were captured using a Chromium Single Cell 3’ Library & Gel Bead Kit (V3; 10× Genomics) according to the manufacturer’s instructions. The cDNA amplification and library construction procedures were performed according to standard protocols. The resulting libraries were sequenced on the Illumina sequencing platform by LC-BIO Co., Ltd (HangZhou, China).

Cell Ranger (Version 6.1.2) was used to process the single-cell data, align the reads, and generate feature-barcode matrices. Homo_sapiens_GRCh38_96 was used as the reference genome. Then the matrices from different samples were aggregated using the ‘aggr’ function.

### Quality control, dimension reduction, clustering, and cell-type identification

Basic data processing and visualization was performed with the Seurat package (Version 4.0.2)(Hao et al., 2021). Briefly, data were loaded using the “Read10x” function and a Seurat object was built. The data were log normalized and scaled. Variable genes were identified by the “FindVariableGenes” function. Next, principal component analysis (PCA) was performed, and the top 20 principal components (PCs) were used for UMAP dimension reduction and clustering (resolutions=0.5). We then performed quality control with the following criteria: (1) Clusters with a high percentage of mitochondrial genes were removed; (2) cells with a total number of expressed genes >800, 4000< total UMI count<120,000, and mitochondria genes < 60% were retained. We pre-processed the data again and removed doublet cells with DoubletFinder (Version 2.0.3). The 22,520 cells that remained were taken as having passed quality control and were used for subsequent analyses.

After quality control, data matrices from different donors were integrated by canonical correlation analysis (CCA)(Stuart et al., 2019). Briefly, the data were integrated using the top 4000 variable genes as integration anchors. Then the first 30 PCs were used for UMAP dimension reduction, to construct a k-nearest neighbor (kNN) graph, and to refine the edge weights between any two cells. The cells were then clustered using the Louvain algorithm for modularity optimization with the resolution parameter set to 0.6. Cell types was identified using the indicated marker genes (see Results section). Marker genes for each cluster were determined with ROC analysis using the “FindAllMarkers” function. Only those with |avg_logFC| > 0.4, min.pct = 0.5, and *p*-value < 0.05 were considered as marker genes. GO analysis of cell type-specific markers were performed with clusterProfiler (Version 4.0.5)(Wu et al., 2021).

### Analysis of coefficient of variation

To observe the effects of aging on germline and somatic cells, we performed age-relevant coefficient of variation analysis, as described in previous studies(S. Wang et al., 2020). We identified highly variable genes (HVGs) using the “FindVariableFeatures” function and selected the top-ranked 10% genes as HVGs (3438 HVGs out of 34,378 genes) for downstream aging-associated transcriptional variation analysis.

For a given cell type, *c*, we defined the cell-paired-distance *d* for HVG *x* between the cells in young (denoted as *i*) and old groups (denoted as *j*) as:

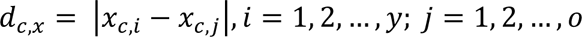

where *y* and *o* are the cell numbers in young and old groups, respectively, of cell type *c*.

Then we calculated the arithmetic mean of *d_c,x_* as 𝜇*_c,x_* and the standard deviation as σ*_c,x_* . Therefore, the coefficient of variation of cell-paired-distance, or transcriptional noise, is defined as:

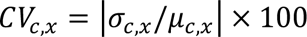

### Identification of differentially expressed genes (DEGs)

The function “FindMarkers” in the R package, Seurat, which is based on the Wilcoxon rank-sum test, was used to identify DEGs. Genes with an average log2-transformed difference greater than 0.25 and *p*-value < 0.05 were considered to be aging-associated DEGs.

### GSEA analysis

Enrichment analysis between young and old groups was performed based on gene set enrichment analysis (GSEA), which was performed using fgsea (Version 1.16.0). The random seed used in the permutations process was set as 123. Genes were ranked according to fold change, and the gene fold-change data frame was used as the input. The GO-BP database was used as the reference database, and was loaded by msigdbr (Version 7.4.1). All other parameters used to perform GSEA were set at the default values. We filtered the obtained GSEA items with *p*-value < 0.05 and |normalized enrichment score (NES)|**>** 1. The results were visualized using ggplot2 (Version 3.3.5).

### Analysis of gene regulation networks

Gene regulation network analysis was performed based on the single-cell regulatory network inference (SCENIC) workflow using default parameters. Firstly, the quality controlled, log-transformed UMI count matrix and TFs were loaded as input. For the UMI count matrix, the DEGs between age groups were depicted as row names, and the cell barcodes of each cell type were represented as column names. Secondly, the correlation matrix of genes was constructed for network inference using the random forest-based algorithm applied by GENIE3 (Version 1.12.0)(Huynh-Thu, Irrthum, Wehenkel, & Geurts, 2010). Reference TFs were downloaded from RcisTarget (https://resources.aertslab.org/cistarget). Thirdly, based on the RcisTarget database, coexpression modules enriched for target genes of each candidate TF were detected by SCENIC (Version 1.2.4)(Aibar et al., 2017). The activity of each TF module in each cell was computed by AUCell (Version 1.12.0), and the regulation networks of TF modules with high normalized enrichment scores were visualized by Cytoscape (Version 3.8.2).

### AUCell

To score individual cells for pathway activities, we used the R package, AUCell (Version 1.12.0). First, a log-normalized expression matrix was used as input to compute gene-expression rankings in each cell by applying the “AUCell_buildRankings” function with default parameters. The genes in each gene set are listed in Supplementary file 4. The activity of canonical pathway gene sets for each cell was then computed by scoring the area under the curve (AUC) values based on gene expression rankings, using the “AUCell_calcAUC” function. The score of each cell was visualized with the R package application, ggplot2 (Version 3.3.5).

### Cell trajectory analysis

Pseudotime trajectory analysis was performed using monocle (Version 2.18.0)(Qiu et al., 2017). The UMI count matrix and meta data were extracted from the quality-controlled Seurat object and loaded to the monocle using the function, “newCellDataSet”. Then the ordering genes detected by function “differentialGeneTest” were used to define the process. “DDRTree” was used to reduce the dimensions to one with two dimensions. Each cell trajectory was then constructed using the “orderCells” function.

### Cell-cell communication analysis

The analysis of cell-cell communication by ligand-receptor pairs was performed using CellChat (Version 1.1.3)(Jin et al., 2021). First, the normalized data matrix and meta data were extracted from the quality-controlled Seurat object and loaded. The “secreting signal” of CellChatDB was used as the ligand-receptor interaction database. Once the overexpressed genes were identified, the between-cell interaction pairs and interaction probabilities for each L-R pair were calculated. Then the communication probability on a signaling pathway level was calculated by summarizing all related ligands/receptors using the function, “computeCommunProbPathway”. Finally, the young and old CellChat objects were merged for comparison.

### Immunofluorescence staining

Human testis tissues were embedded in paraffin and sectioned at 4 µm. After being deparaffinized by xylene and rehydrated with an ethanol series (100%, 95%, 85%, 75%) at room temperature, the sections were incubated in citrate antigen-retrieval solution (Beyotime, Shanghai, China) in a hot water bath (96℃) for 20 min. For intracellular protein detection, the sections were permeabilized with 0.5% Triton X-100 (Sigma-Aldrich, St. Louis, MO, USA) for 20 min. The tissue sections were then incubated with 3% BSA for 30 min at room temperature and incubated with primary antibodies overnight at 4℃. Following incubation, the tissue sections were rinsed five times with PBS and incubated with secondary antibodies for 60 min at room temperature. After being rinsed five times with PBS, the tissue sections were stained with 4,6-diamidino-2-phenylindole (DAPI; Invitrogen, Carlsbad, CA, USA) for 5 min, after which the DAPI was removed and replaced with mounting medium (DAKO; Glostrup, Denmark). The specific fluorescence was visualized and photographed using an LSM800 confocal microscope (Zeiss, Jena, Germany). The utilized primary and secondary antibodies are listed in Supplementary file 7.

### Isolation and culture of LCs

Primary human LCs were isolated from human testicular tissues as previously described(Luo et al., 2021). In brief, cryopreserved testicular tissues were thawed quickly and washed three times with PBS. Then, the testes were mechanically cut and enzymatically dissociated using 1 mg/mL type IV collagenase and 200 μg/mL DNase I dissolved in DMEM/F-12 for 20 min with slow shaking (100 cycles/min) at 37℃. The samples were filtered through a 40-μm filter and centrifuged at 300g for 4 min. The cell pellets were rinsed twice with PBS and resuspended in PBS containing 0.1% BSA for fluorescence-activated cell sorting (FACS) (MoFlo Astrios EQs, Beckman Coulter, CA, USA). Before LC isolation, we used a two-step gating strategy to eliminate debris and cell doublets based on the side scatter (SSC) and forward scatter (FSC) parameters. Then, the cell samples were analyzed in the combined fluorescence channels (405- 448 and 640-671 nm) of flow cytometry, and the LC population distinguished from the main group was isolated.

The obtained primary LCs were cultured in medium containing DMEM/F12, 10% FBS and 1% insulin-transferrin-sodium selenite (ITS; Thermo Fisher Scientific) at 35℃ with 5% CO2. The ability of the cells to produce testosterone was assessed after 3 h of incubation with DMEM/F12 containing 0.1% BSA, 1 IU/mL hCG (R&D, Systems, Minneapolis, USA), 10 μM 22-HC (Sigma-Aldrich), and 1× ITS. The cell supernatants were collected and stored at −80℃ until analysis.

### Ex vivo culture of testicular tissues

For short-term tissue culture, we modified a previously described method(X. Li et al., 2016). Briefly, cryopreserved testicular tissues were thawed quickly and washed three times with PBS. Then, the testes were mechanically cut into small pieces (2∼4 mm^2^ in size). Three pieces of tissues were plated per well of a 12-well plate and cultured in medium containing DMEM/F12, 0.1% BSA, and 1× ITS (Gibco) at 35℃ with 5% CO2.

The ability of the tissues to produce testosterone was assessed after 3 h of incubation with DMEM/F12 (Gibco) containing 0.1% BSA, 1 IU/mL hCG (R&D), 10 μM 22-HC (Sigma-Aldrich), and 1× ITS (Gibco). The supernatants to be used for the testosterone assay were collected and stored at −80℃ until analysis. Tissues were collected and lysed in cold RIPA buffer, followed by protein quantification for statistical analysis.

### Antioxidant treatments

To elucidate the protective effect of antioxidants on cellular oxidative stress and testosterone production, primary LCs or testicular tissues from the old group were cultured with DMSO, 10 mM N-acetyl-L-cysteine (NAC; MedChemExpress, Shanghai, China), and 50 μM vitamin E (MedChemExpress). After a 24 h culture period, LCs or tissues were used for subsequent analysis.

### Testosterone measurements

Testosterone concentrations were assayed as previously reported by our group(Luo et al., 2021). The cell or tissue supernatants were collected at the indicated timepoints and stored at −80℃ until analysis. Testosterone levels were measured using a chemiluminescent immunoassay (CLIA) system (Architect system; Abbott GmbH & Co. KG, Germany). The coefficient of variation of this CLIA system is 1.9–5.1% for intra-assay precision and 2.5–5.2% for inter-assay precision. The lowest detectable dose of testosterone was 0.01 ng/mL.

### ROS detection

Intracellular ROS was detected using a Reactive Oxygen Species Assay Kit (Thermo Fisher Scientific) according to the manufacturer’s instructions. Briefly, primary LCs were collected and incubated with DMEM/F12 medium containing 1 μM DHE for 60 min at 37℃ in the dark, and fluorescent intensity (Ex = 495 nm, Em = 520 nm) was measured by flow cytometry (CytoFLEX; Beckman Coulter, CA, USA) or photographed under a DMi8 microscope (Leica).

### Statistical analysis

All data were analyzed using GraphPad Prism v8 (GraphPad Software, La Jolla, CA, USA). Statistical differences between samples were assessed with Student’s t-tests, one-way analysis of variance (ANOVA). Differences were considered significant when *p* < 0.05 (\**p* < 0.05, \*\**p* < 0.01 and \*\*\**p* < 0.001).

## Acknowledgements

We thank the patients and their families for their dedication. We thank Dr. Fangjian Zhou and Dr. Yonghong Li for coordinating the human testicular tissue collection; We are grateful to Yan Guo, Hong Chen, Xiangqian Guo and Xikun Han for their constructive suggestions.

## Additional information

### Funding

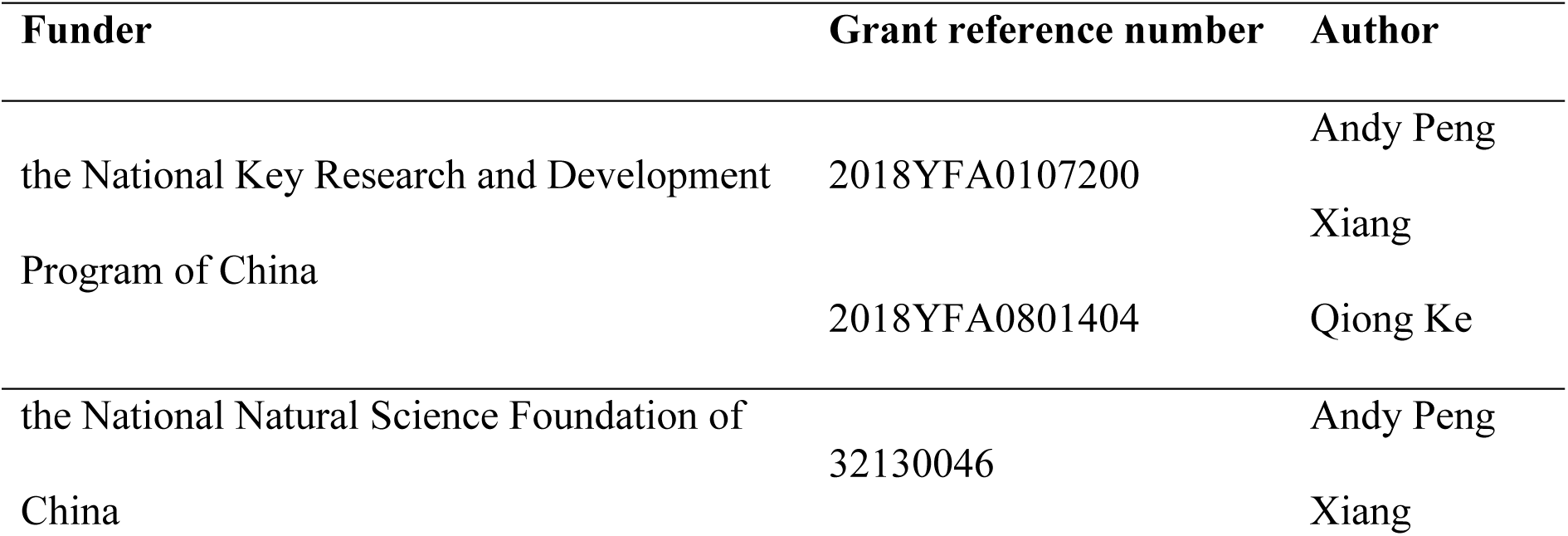
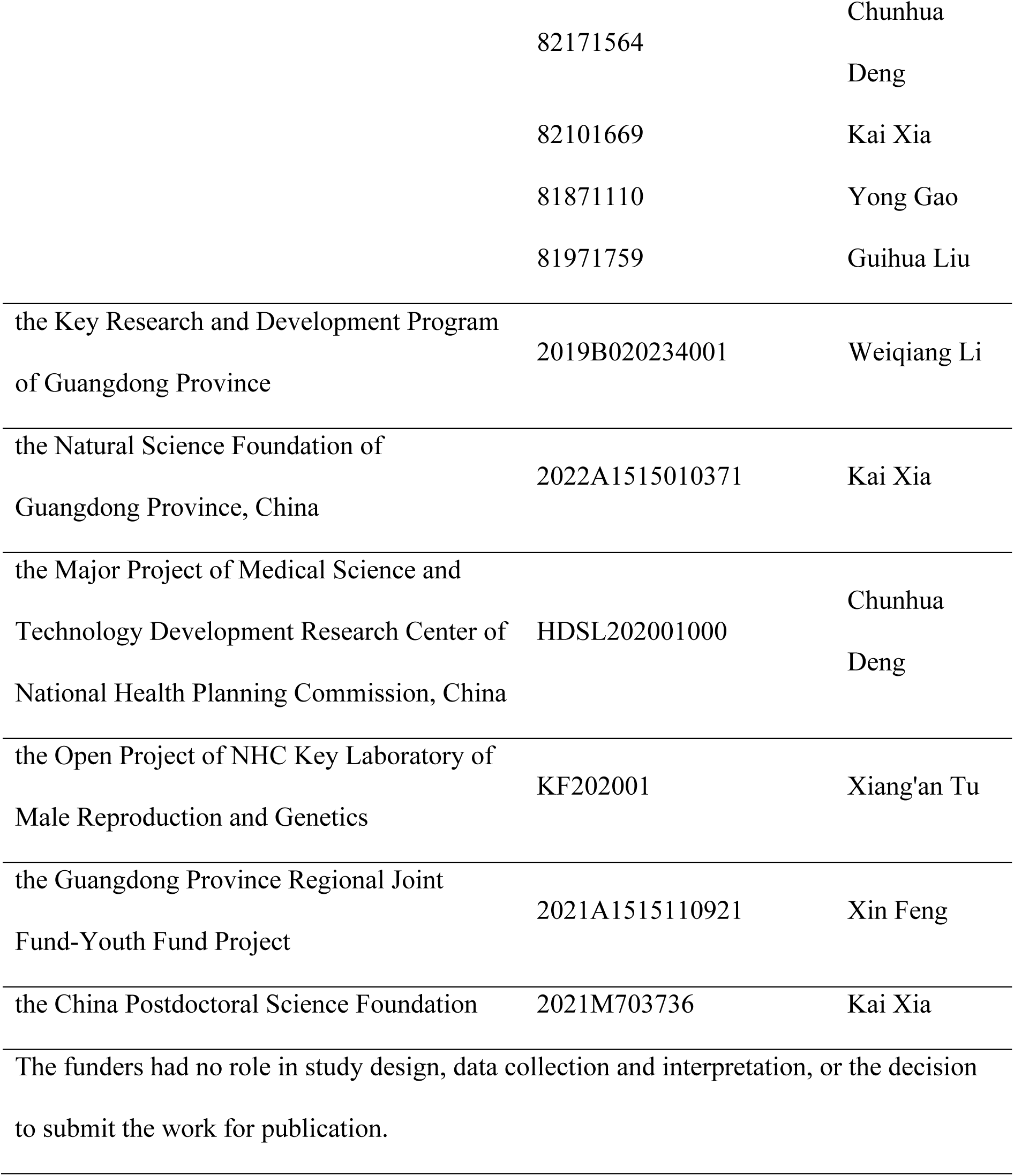

### Author contributions

Kai Xia, Conceptualization, Methodology, Validation, Formal analysis, Data curation, Writing – original draft preparation, Funding acquisition; Siyuan He, Methodology, Formal analysis, Investigation, Visualization; Peng Luo, Validation, Investigation, Resources; Lin Dong, Validation, Investigation, Visualization; Feng Gao, Validation, Investigation; Xuren Chen, Validation, Investigation; Yunlin Ye, Investigation, Resources; Yong Gao, Investigation, Resources, Funding acquisition; Yuanchen Ma, Data curation; Yadong Zhang, Data curation, Visualization, Funding acquisition; Qiyun Yang, Investigation, Resources; Dayu Han, Investigation, Resources; Xin Feng, Investigation, Resources, Funding acquisition; Zi Wan, Investigation, Resources; Hongcai Cai, Investigation, Resources; Qiong Ke, Investigation, Resources, Funding acquisition; Tao Wang, Investigation, Resources; Weiqiang Li, Resources, Funding acquisition; Xiang’an Tu, Resources, Funding acquisition; Xiangzhou Sun, Resources; Chunhua Deng, Conceptualization, Resources, Writing – review & editing, Supervision, Project administration, Funding acquisition; Andy Peng Xiang, Conceptualization, Resources, Writing – review & editing, Supervision, Project administration, Funding acquisition.

### Competing interests

The authors declare no competing interests.

### Ethics

The protocols were approved by the Ethics Committee of the First Affiliated Hospital of Sun Yat-sen University (Assurance # 2019-148).

## Additional files

### Supplementary files

Supplementary file 1. Cell cluster information, markers, GO terms, and transcriptional regulators for each cell type in human testis; related to Figure 1 and Figure 1-Figure supplement 1.

Supplementary file 2. Differential gene expression analysis results; related to Figures 3, 4, Figure 4-Figure supplement 1, and 5.

Supplementary file 3. GSEA analysis of testicular cells; related to Figures 3, Figure 3-Figure supplement 1, 4, and 5.

Supplementary file 4. Gene lists for base excision repair (BER), nucleotide excision repair (NER), and senescence-associated secretory phenotype (SASP); related to Figures 3 and 4.

Supplementary file 5. List of LCs regulons obtained from the SCENIC analysis; related to Figure 5.

Supplementary file 6. Ligand-receptor pairs of testicular cells obtained from CellChat analysis; related to Figures 6 and Figures 6-Figure supplement 1.

Supplementary file 7. The antibodies used in this study; related to the Methods.

### Source data files

Figure 1–source data 1, Related to Figure 1A–B

Figure 2–source data 1, Related to Figure 2F–H

Figure 3–source data 1, Related to Figure 3F–G

Figure 4–Figure supplement 1–source data 1, Related to Figure 4–Figure supplement 1A–C

Figure 5–source data 1, Related to Figure 5F–J

Figure 5–Figure supplement 1–source data 1, Related to Figure 5–Figure supplement 1B–C

## Data availability

The RNA-seq sequencing and processed data reported in this paper have been deposited in the Genome Sequence Archive (GSA for Human) with project number HRA002349.

The following dataset was generated:

| Author(s) | Year | Dataset title | Dataset URL | Database and Identifier |
| --- | --- | --- | --- | --- |
| Kai Xia | 2022 | Single-Cell | https://ngdc.cnc |  |
|  |  | Transcriptomic | [b.ac.cn/search/?](https://b.ac.cn/search/?dbId=hra&q=HRA002349) | GSA for Human, |
|  |  | Landscape of Human | [dbId=hra&q=H](https://b.ac.cn/search/?dbId=hra&q=HRA002349) | HRA002349 |
|  |  | Testicular Aging | [RA002349](https://b.ac.cn/search/?dbId=hra&q=HRA002349) |  |

---

**Figure 1 supplement 1.**
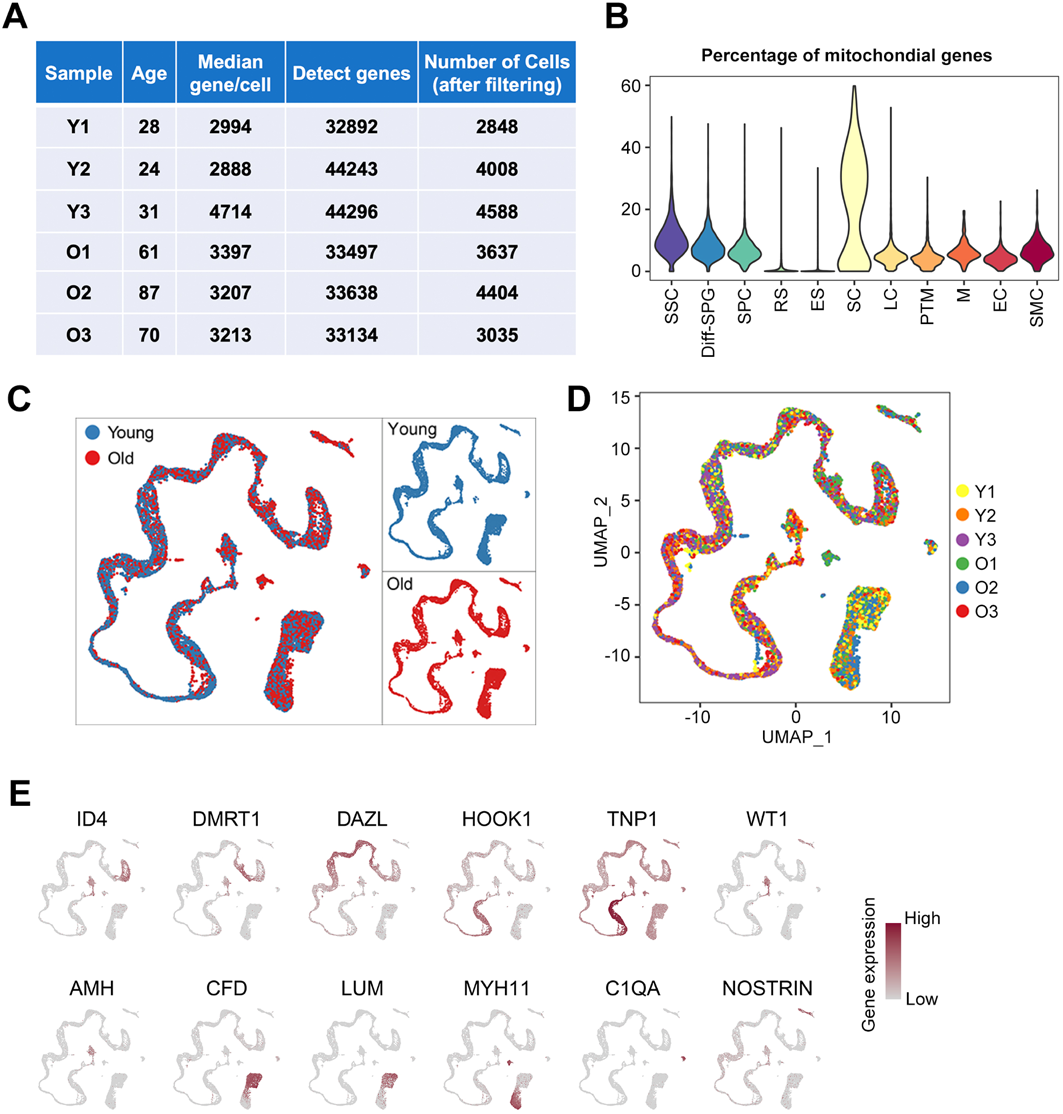
Information on Human Samples and Quality Control of Single-Cell RNA-Seq. (A) Summary information for the testicular samples analyzed in this study. (B) Violin plot showing the percentages of mitochondrial genes detected in each cell type. (C) UMAP plot showing the cell distributions for the young (red) and old (blue) groups. (D) UMAP plot showing the cell distributions for each sample. Cells are colored and annotated to the right. (E) UMAP plot showing the expression profiles of cell-type-specific marker genes for the indicated cell types in human testis. The color key, ranging from gray to brown, indicates low to high gene expression levels.

**Figure 1 supplement 2.**
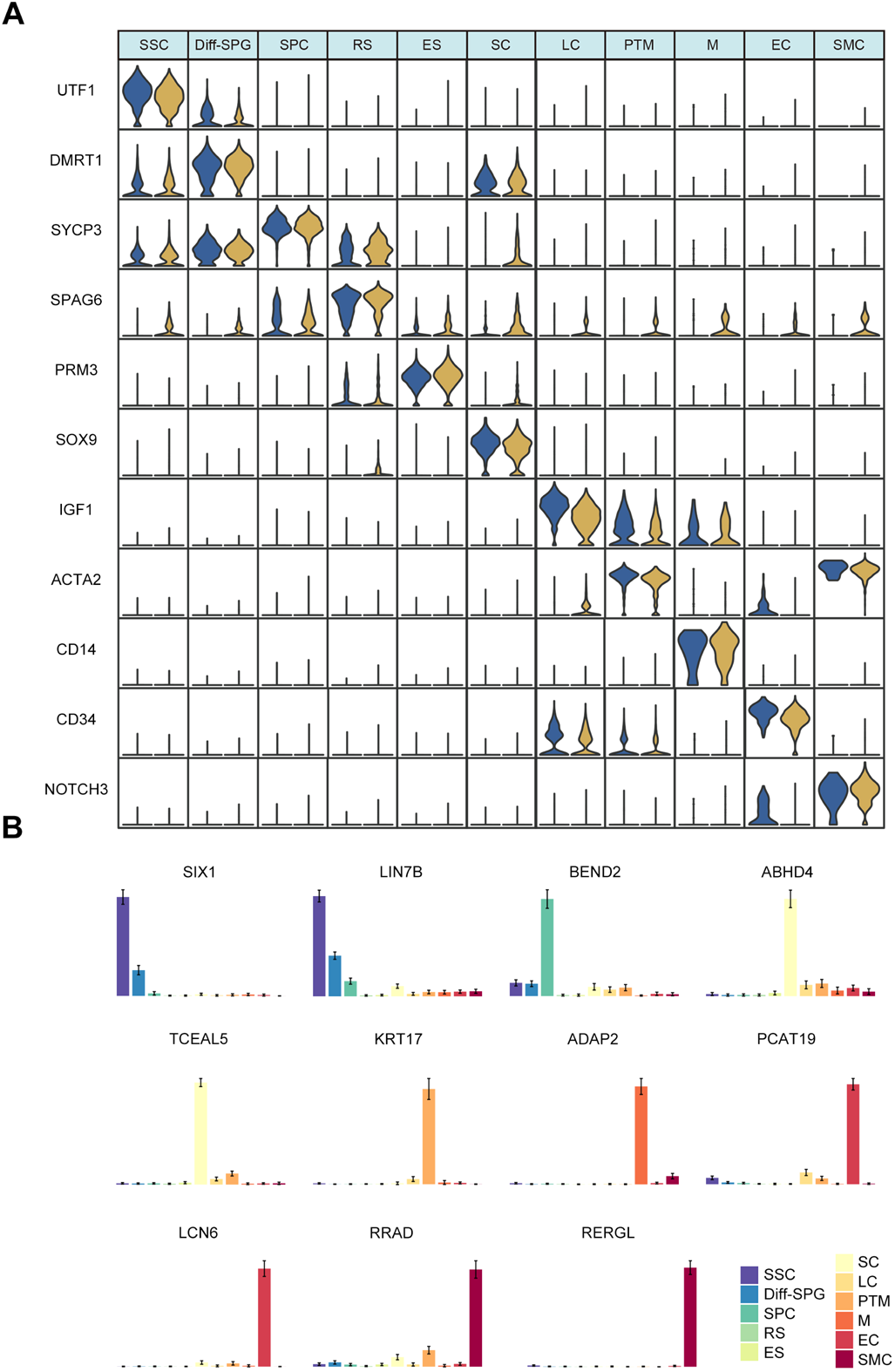
The Expression Level of Marker Genes for Each Cell Type. (A) Violin plot showing expression levels of marker genes for each cell type in young and old human testis. (B) Bar plot showing the expression levels of representative new marker genes for various cell types.

**Figure 3 supplement 1.**
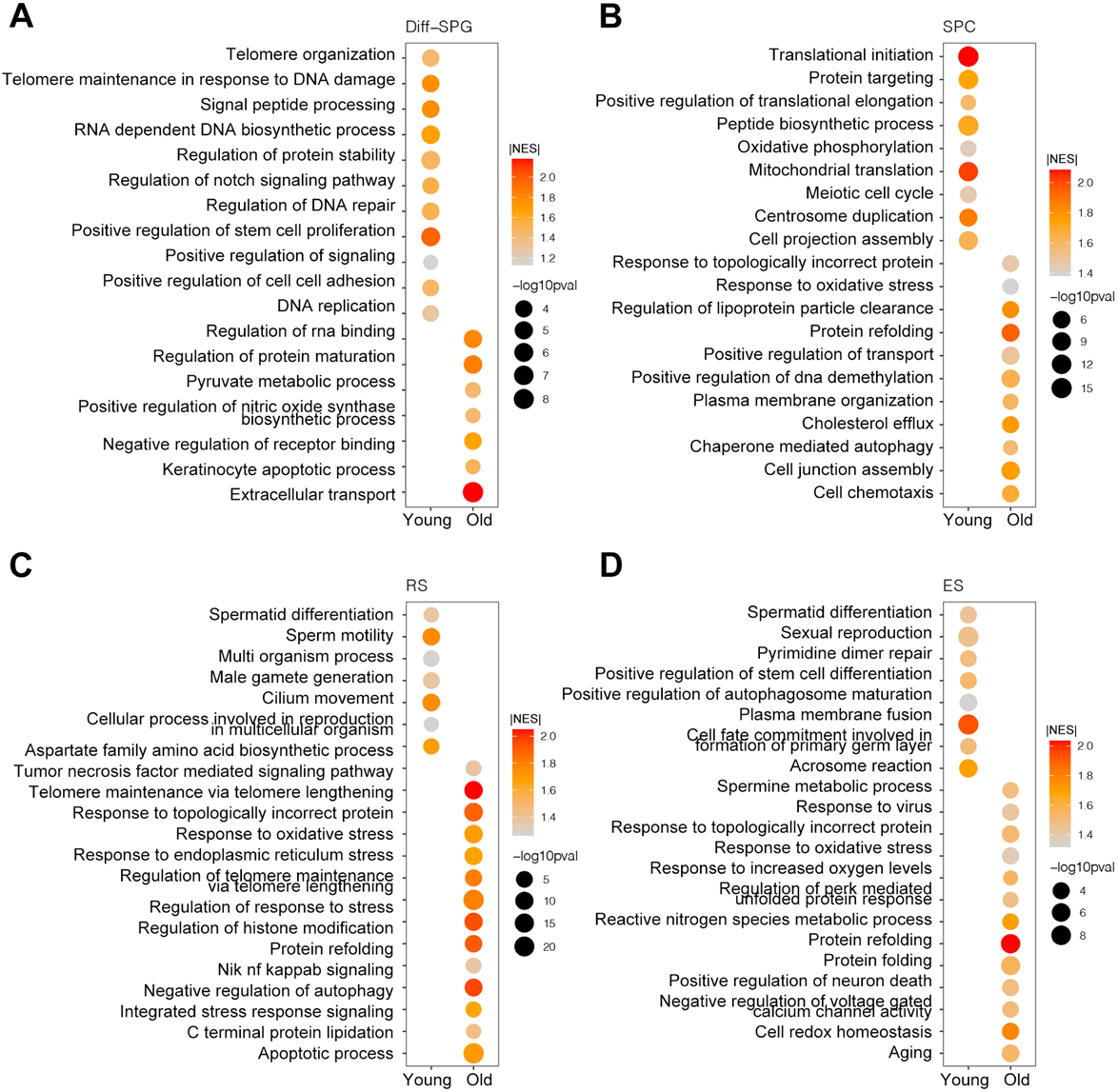
GO Analysis Between the Different Germ Cell Clusters of the Young and Old Datasets. (A-D) Representative Aging-Associated GO terms of DEGs in Diff-SPGs (A), SPCs (B), RSs (C), and ESs (D). Dot size indicates the range of *p*-value. The color keys, ranging from gray to red (top) or gray to blue (bottom), indicate the absolute value of the normalized enrichment score (NES).

**Figure 4 supplement 1.**
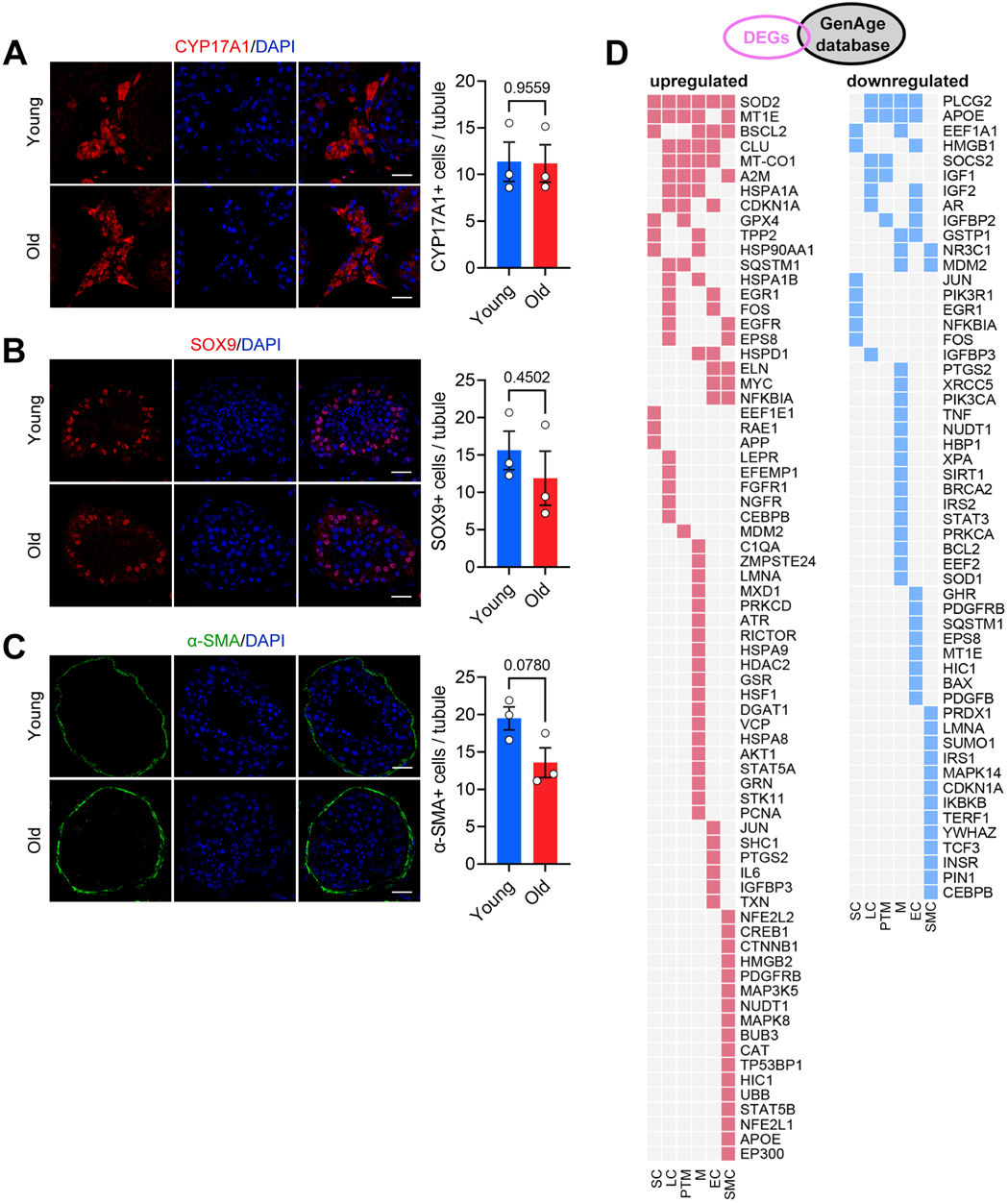
Changes in the Cellular and Transcriptional Regulatory Networks of Somatic Cell Types during Human Testicular Aging. (A-C) Immunostaining and quantification of CYP17A1 (A), SOX9 (B), and α-SMA (C) in young and old human testis sections. Left, representative image of CYP17A1 (A), SOX9 (B), and α-SMA (C) in human testis sections. Right, quantification of CYP17A1+ (A), SOX9 (B)+, and α-SMA+ (C) cells per seminiferous tubule. Scale bars, 35 μm. Young, n=3; Old, n=3. Data are expressed as mean ± sem. Significance was determined by Student’s t-test. (D) Heatmap showing expression patterns of upregulated (red) and downregulated (blue) genes found in the GenAge database.

**Figure 5 supplement 1.**
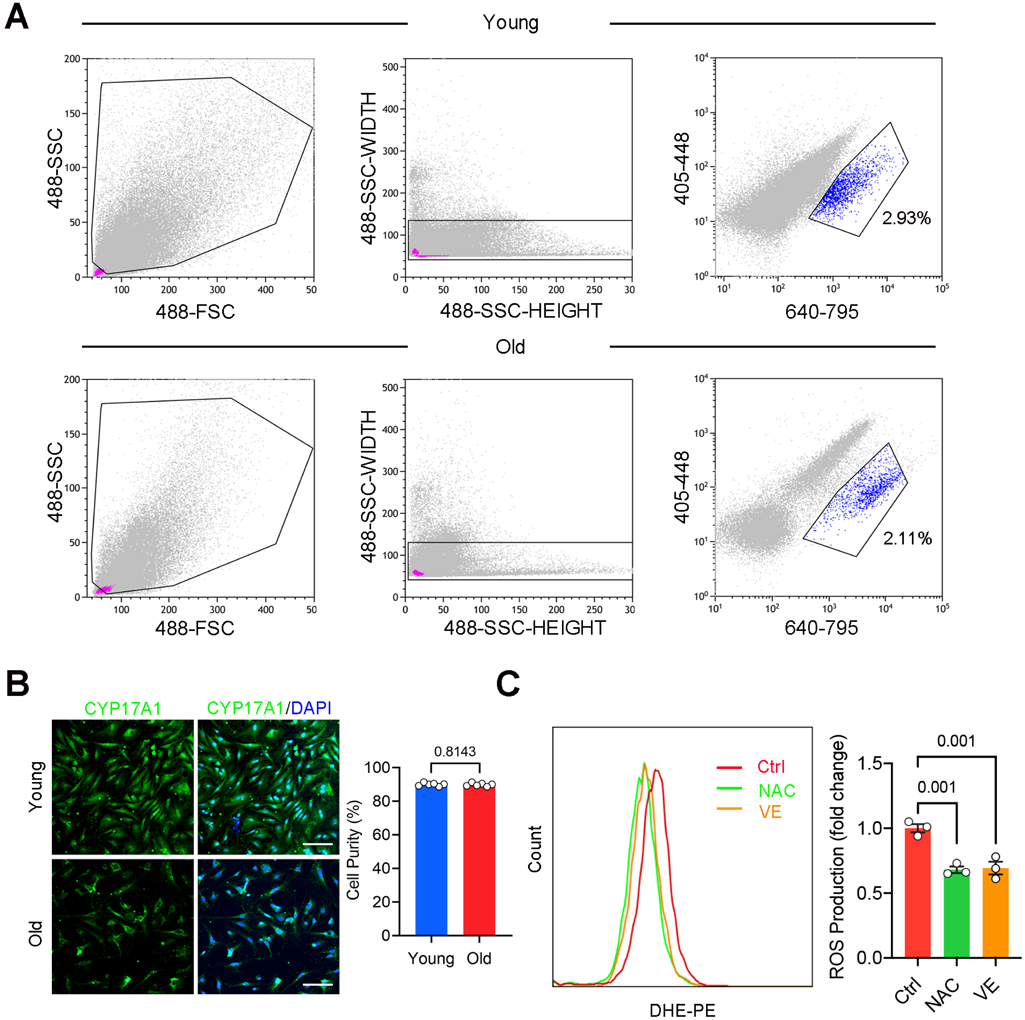
Isolation and ROS quantification of primary human LCs. (A) Representative gating strategy of FACS for isolating young and old human LCs. Autofluorescent cells were excited in all fluorescence channels and isolated with the 405/448 merged 640/795 channel. FSC: forward scatter; SSC: side scatter. Cell purity of young and old human LCs was measured by immunostaining of CYP17A1. Right, quantification of the proportion of CYP17A1+ cells. Scale bars, 100 μm. Young, n=6; Old, n=6. Data are expressed as mean ± sem. Significance was determined by Student’s t-test. (C) Quantification of ROS production in primary LCs of control and antioxidant-treated (NAC or VE) groups, as measured by flow cytometry. Control, n=3; NAC (10 mM), n=3; VE (5 μM), n=3. Data are expressed as mean ± sem. Significance was determined by one-way ANOVA.

**Figure 6 supplement 1.**
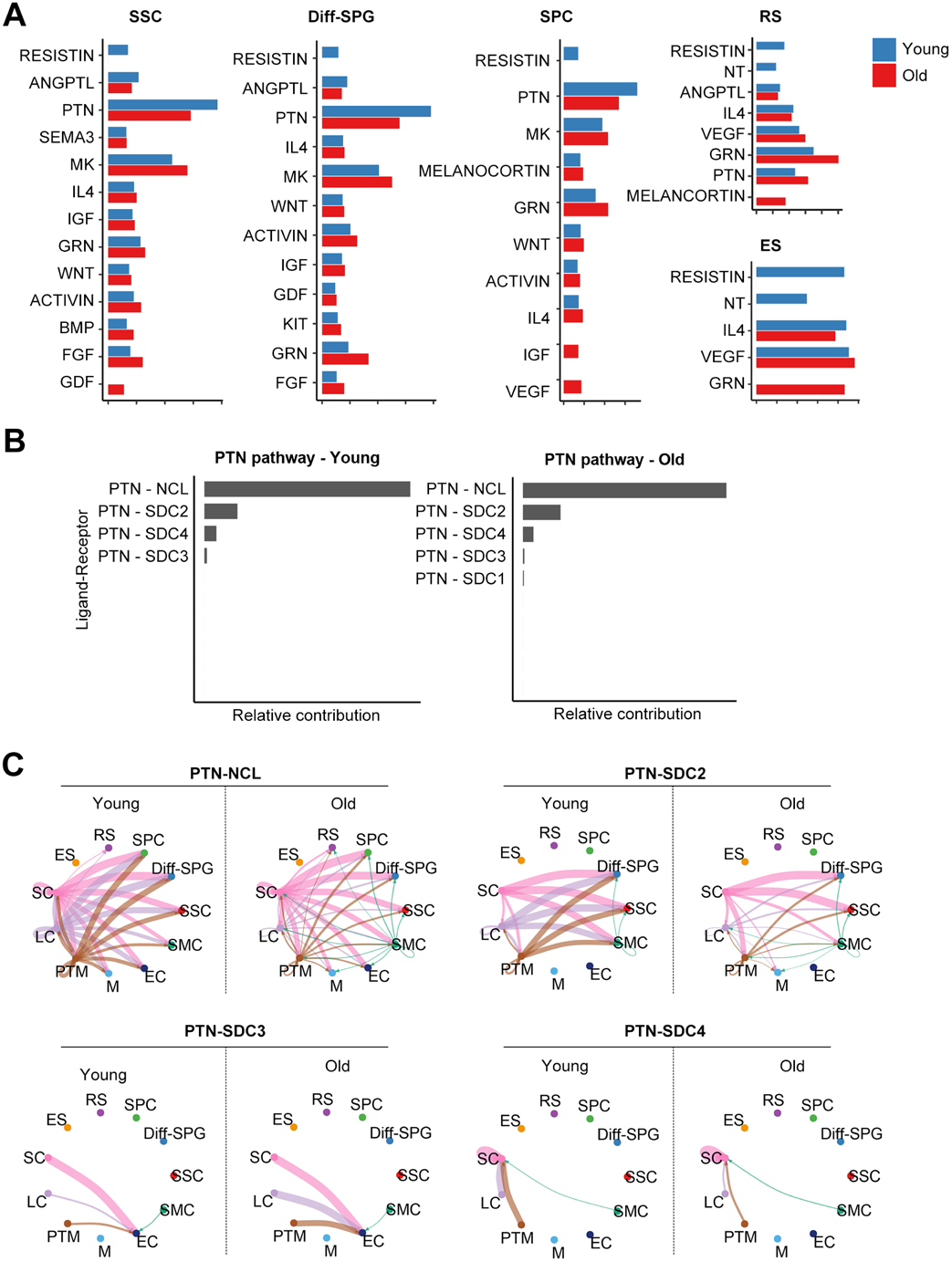
PTN signaling pathways changes during human testicular aging. (A) Information flow of significant signaling pathways received by germ cell populations in the young (red columns) and old (blue columns) groups. PTN signaling pathway is labeled in red. (B) Relative contribution of each ligand-receptor pair to the overall PTN signaling network in young (left) and old (right) groups. In the PTN signaling pathway, PTN acts as a ligand for the receptors, NCL, SDC2, SDC3, and SDC4. (C) Circle plot showing the interaction probability between all cell populations of each ligand-receptor pair in the PTN signaling pathway. Edge width represents the communication probability. In the PTN signaling pathway, PTN acts as a ligand for the receptors, NCL, SDC2, SDC3, and SDC4.

## Notes

### Competing Interest Statement

The authors have declared no competing interest.

